# GRK2 Kinases in the Primary Cilium Initiate SMOOTHENED-PKA Signaling in the Hedgehog Cascade

**DOI:** 10.1101/2023.05.10.540226

**Authors:** Madison F. Walker, Jingyi Zhang, William Steiner, Pei-I Ku, Ju-Fen Zhu, Zachary Michaelson, Yu-Chen Yen, Annabel Lee, Alyssa B. Long, Mattie J. Casey, Abhishek Poddar, Isaac B. Nelson, Corvin D. Arveseth, Falko Nagel, Ryan Clough, Sarah LaPotin, Kristen M. Kwan, Stefan Schulz, Rodney A. Stewart, John J. G. Tesmer, Tamara Caspary, Radhika Subramanian, Xuecai Ge, Benjamin R. Myers

## Abstract

During Hedgehog (Hh) signal transduction in development and disease, the atypical G protein-coupled receptor (GPCR) SMOOTHENED (SMO) communicates with GLI transcription factors by binding the protein kinase A catalytic subunit (PKA-C) and physically blocking its enzymatic activity. Here we show that GPCR kinase 2 (GRK2) orchestrates this process during endogenous Hh pathway activation in the vertebrate primary cilium. Upon SMO activation, GRK2 rapidly relocalizes from the ciliary base to the shaft, triggering SMO phosphorylation and PKA-C interaction. Reconstitution studies reveal that GRK2 phosphorylation enables active SMO to bind PKA-C directly. Lastly, the SMO-GRK2-PKA pathway underlies Hh signal transduction in a range of cellular and *in vivo* models. Thus, GRK2 phosphorylation of ciliary SMO, and the ensuing PKA-C binding and inactivation, are critical initiating events for the intracellular steps in Hh signaling. More broadly, our study suggests an expanded role for GRKs in enabling direct GPCR interactions with diverse intracellular effectors.

## INTRODUCTION

The Hedgehog (Hh) signaling pathway is a cornerstone of animal development, homeostasis, and disease, controlling the formation of nearly every vertebrate organ and instigating several common malignancies^1–6^. To carry out these roles, the Hh pathway utilizes an intracellular transduction cascade that originates at the cell surface and propagates to the nucleus, culminating in transcription of genes linked to proliferation or differentiation^1–4^. This process relies on the primary cilium, a tiny antenna-shaped cell surface compartment thought to facilitate the underlying biochemical steps via its highly confined ultrastructure and its unique protein and lipid inventory^7–12^. Despite extensive research, the mechanisms by which Hh signals are transmitted from the cell surface to the nucleus remain longstanding mysteries^1–4,12^.

In the Hh pathway “off” state, the atypical G protein-coupled receptor (GPCR) SMOOTHENED (SMO) is inhibited within the cilium by PATCHED1 (PTCH1), a 12-transmembrane (TM) sterol transporter that restricts access of SMO to endogenous activating sterol ligands^2–4^. Consequently, SMO is rapidly cleared from the cilium via ubiquitylation and engagement of the ciliary exit machinery^13–17^, and the protein kinase A catalytic subunit (PKA-C) is free to phosphorylate and inactivate GLI transcription factors^18–25^. In the pathway “on” state, Hh proteins bind to and inhibit PTCH1, enabling SMO to bind sterols^26–29^, assume an active conformation^26–29^, and accumulate to high levels in the cilium^30–32^. Active SMO blocks PKA-C phosphorylation of GLI, leading to GLI activation^22,24,25,33,34^. SMO inhibition of PKA-C is thus fundamental to Hh signal transduction, but the underlying mechanism has remained poorly understood, mainly because it does not rely on the standard signaling paradigms employed by nearly all other GPCRs^35–37^.

We recently discovered that SMO can inactivate PKA-C via an unusual mechanism: it utilizes a decoy substrate sequence, termed the “protein kinase inhibitor” (PKI) motif, to directly bind the PKA-C active site and physically block its enzymatic activity. SMO can bind and inhibit PKA-C as a decoy substrate *in vitro*, and these decoy substrate interactions are necessary for Hh signal transduction in cultured cells and *in vivo.* Based on these findings, we proposed that SMO utilizes its PKI motif to inactivate PKA-C in cilia, thereby blocking GLI phosphorylation and triggering GLI activation^38,39^.

The above model immediately raises a fundamental question: what controls SMO / PKA-C interactions such that they occur only when SMO is in an active state, an essential condition for faithful Hh signal transmission and for avoiding pathological outcomes resulting from insufficient or excessive GLI activation? Indeed, we found that during Hh signal transduction in cilia, only the active, agonist-bound conformation of SMO is competent to colocalize with PKA-C^39^. To explain this phenomenon, we proposed that GPCR kinase (GRK) 2 (along with its paralog, GRK3, hereafter referred to collectively as “GRK2”) regulate SMO / PKA-C interactions^38,39^. GRKs are a multigene family that bind selectively to the active conformations of GPCRs and phosphorylate their intracellular domains, triggering interactions between GPCRs and other signaling proteins^40–42^. GRK2 is essential for SMO-GLI communication^43–45^, and this kinase selectively phosphorylates active, agonist-bound SMO expressed in HEK293 cells^38,46^, thereby enhancing interactions between the SMO PKI motif and PKA-C^38,39^. Furthermore, blocking GRK2 kinase activity^43,44^ or mutating the GRK2 phosphorylation sites on SMO^38^, which we mapped via mass spectrometry^38^, reduces SMO / PKA-C interactions in HEK293 cells^38^ and blocks Hh signal transduction *in vivo*^43,44^.

Although these findings suggest that GRK2 can serve as an intermediary between SMO and PKA-C, they leave the physiological role of the SMO-GRK2-PKA communication pathway, as well as its underlying molecular mechanism, unresolved in several respects. First, whether SMO undergoes GRK2-mediated phosphorylation during physiological Hh signal transduction in the primary cilium is not known. Whereas all other essential Hh pathway components localize in or near the cilium^20,30–32,47^, whether this is true for GRK2 remains unknown. In addition, our strategy to identify GRK2 phosphorylation sites on SMO has limitations, as it involved overexpression of a truncated, stabilized SMO in a non-ciliated cell line, and utilized biochemical detection methods that lack subcellular resolution^38^. Consequently, GRK2 phosphorylation of SMO, and its role in SMO / PKA-C interactions, have not been studied during endogenous Hh signal transduction on physiologically relevant time scales in primary cilia. Second, while GRK2 phosphorylation contributes to formation of SMO / PKA-C complexes^38^, it remains unclear whether phosphorylation directly triggers SMO to bind PKA-C, or whether the effect of GRK2 phosphorylation requires additional proteins. This is a critical question because GRK2 phosphorylation of GPCRs is canonically associated with binding of β-arrestins (which serve as scaffolds for a variety of cytoplasmic signaling proteins)^40–42^, and has not been previously demonstrated to induce direct interactions between a GPCR and PKA-C. Finally, our new model for SMO / PKA-C communication has only been evaluated in cultured fibroblasts and zebrafish somitic muscle *in vivo*^38,39^. Thus, we do not yet know whether the SMO-GRK2-PKA pathway is a general aspect of Hh signaling beyond these specific systems.

Here we address the molecular mechanism of SMO-GLI communication by developing novel imaging modalities and immunological detection reagents to monitor the SMO-GRK2-PKA pathway in physiological models of Hh signal transduction, along with reconstitution of pathway components. Using these approaches, we show that SMO, upon activation, undergoes rapid phosphorylation by a pool of GRK2 recruited from the base to the shaft of the cilium, enabling formation of ciliary SMO / PKA-C complexes. We also find that GRK2 phosphorylation is sufficient to trigger direct SMO / PKA-C interactions in a purified *in vitro* system with no other proteins present. Finally, we demonstrate that SMO undergoes GRK2-mediated phosphorylation during Hh signal transduction in multiple tissue, organ, and animal settings. Our work establishes GRK2 phosphorylation of ciliary SMO, and the ensuing PKA-C recruitment, as critical initiating events for transmission of Hh signals from the cell surface to the nucleus. Furthermore, our work provides a blueprint to study the cell biological basis for SMO-PKA communication and other GRK-dependent signaling cascades throughout development, physiology, and disease.

## RESULTS

### GRK2 relocalizes from the base to the shaft of the cilium upon SMO activation

If SMO undergoes GRK2 phosphorylation during Hh signal transduction, GRK2 should be detectable in the cilium. Although a prior study detected endogenous GRK2 protein at the centrosome^48^, whether GRK2 can also localize to the cilium remains unknown. Initial attempts to visualize GRK2 in the cilium via immunofluorescence were confounded by an abundant GRK2 signal in the cell body (data not shown). Because kinase-substrate interactions are typically fleeting and dynamic^49,50^, we suspected that sensitive imaging methods capable of tracking GRK2 localization over extended time periods might reveal patterns of transient, low-level ciliary localization over the high GRK2 signal emanating from the cell body.

We therefore established a live-cilium total internal reflection fluorescence (TIRF) microscopy method to study dynamic protein localization to the cilium over long (multi-hour) time intervals (Fig. 1A). We stably expressed a GRK2-eGFP fusion in inner medullary collecting duct (IMCD3) cells (Fig. S1A, see also “Methods”), a well-established cultured cell model for ciliary imaging studies^38,51,52^, and labeled cilia with a live-cell tubulin dye (SiR Tubulin) (Fig. 1B) To select cells for TIRF studies, eGFP-positive cells were imaged for 15 minutes prior to addition of SAG21k^53^ (a high-affinity derivative of the widely utilized SMO agonist SAG^54^). Only cells exhibiting stable ciliary attachment to the coverslip surface during this period were imaged and analyzed further (n=150). TIRF imaging revealed a punctate GRK2-eGFP signal at the ciliary base in 142 cells (∼95%) prior to SAG21k treatment (Fig. 1C-D, Fig. S1-S3, Supplementary Movie 1). Following addition of SAG21k, a faint but detectable eGFP signal appeared along the entire ciliary shaft in approximately 53% (79/150) of cells, consistent with GRK2 recruitment to activated SMO within the cilium (Fig. 1C-D, Fig. S1, Supplementary Movie 1). GRK2-eGFP was detectable as early as 3 minutes following SAG21k treatment in some cases, although the kinetics varied among cilia (from 3 to 105 min, median = 21 min) (Fig. 1E, Fig. S1B-C). We suspect that the variability arises due to challenges in detecting the low eGFP signal in the cilium relative to the cell body, rather than intrinsic variability in the kinetics of GRK2 ciliary localization. Once GRK2-eGFP appeared in cilia, it persisted for several hours (the duration of the imaging experiment) (Fig. 1C), indicating that SMO activation elicits a sustained, steady-state redistribution of GRK2 from the base to the shaft of the cilium.

**Figure 1:**
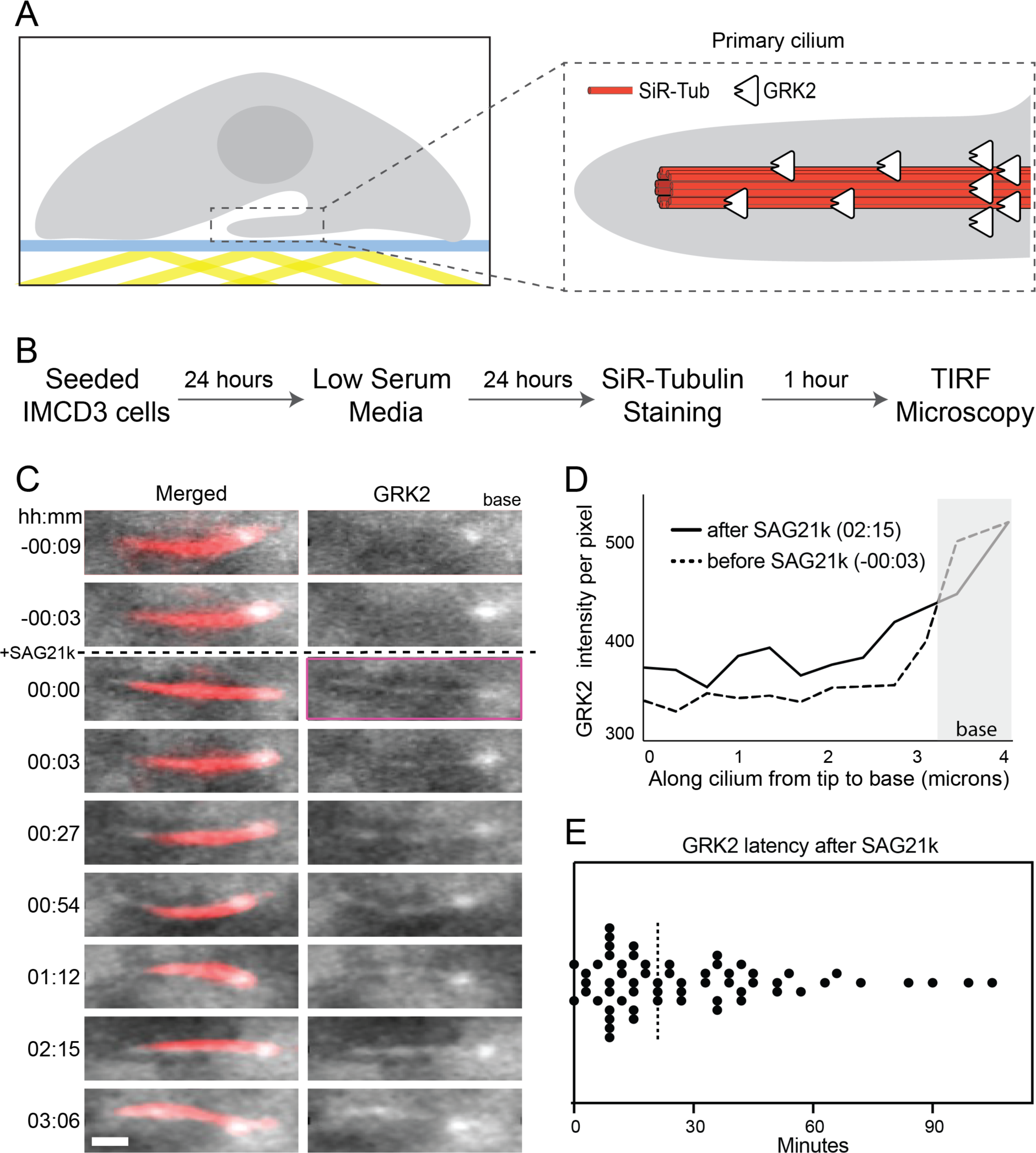
GRK2 relocalizes from the base to the shaft of the cilium upon SMO activation. **(A)** Schematic illustration of a cell with the cilium attached to the coverslip surface for TIRF imaging (left). A close-up view of the cilium (right) where the microtubule was labeled with SiR-Tub and GRK2-eGFP was stably expressed. **(B)** Flowchart outlining the sample preparation procedures for imaging. **(C)** Representative montage from live-cell imaging of a cilium. Merged (red = SiR-Tubulin and white = GRK2-eGFP) and GRK2-eGFP channels are shown. SMO was activated at t = 00:00 (hh:mm) by adding SAG21k. The montages show that eGFP signal was observed in the base of the cilium. After SAG21k addition, eGFP signal is observed in the shaft, and the signal persists for the duration of imaging. The purple box on the GRK2 channel indicates the earliest timepoint following SAG21k addition at which GRK2-eGFP is observed in the ciliary shaft. Scale bar = 1 μm. **(D)** Profiles of GRK2-eGFP intensity per unit area before and after SAG21k activation, for the cilium in (C). **(E)** Scatterplot of the GRK2 latency, defined as the time after SAG21k addition when a GRK2-eGFP signal was observed in the cilium; dashed line indicates median.

Thus, our cilium TIRF imaging studies reveal a pool of GRK2 that localizes to the ciliary base and rapidly redistributes into the ciliary shaft upon SMO activation. These findings are consistent with GRK2 recognition of the SMO active conformation as an early event in intracellular transmission of Hh signals.

### Hh pathway activation leads to accumulation of GRK2-phosphorylated SMO in the cilium

Our GRK2 localization studies suggest that SMO undergoes GRK2-mediated phosphorylation within the cilium. To detect these phosphorylation events directly, we raised a phospho-specific antibody (hereafter referred to as “anti-pSMO”) by immunizing rabbits with a phosphorylated peptide spanning a cluster of highly conserved GRK2 sites that we previously identified in the SMO intracellular membrane-proximal cytoplasmic tail (pCT) (S594, T597, and S599)^38^ (Fig. 2A). Characterization of our anti-pSMO antibody in HEK293 cells transfected with SMO and GRK2 demonstrated that it specifically recognizes the active, GRK2-phosphorylated form of SMO (Fig. S4A-C).

**Figure 2:**
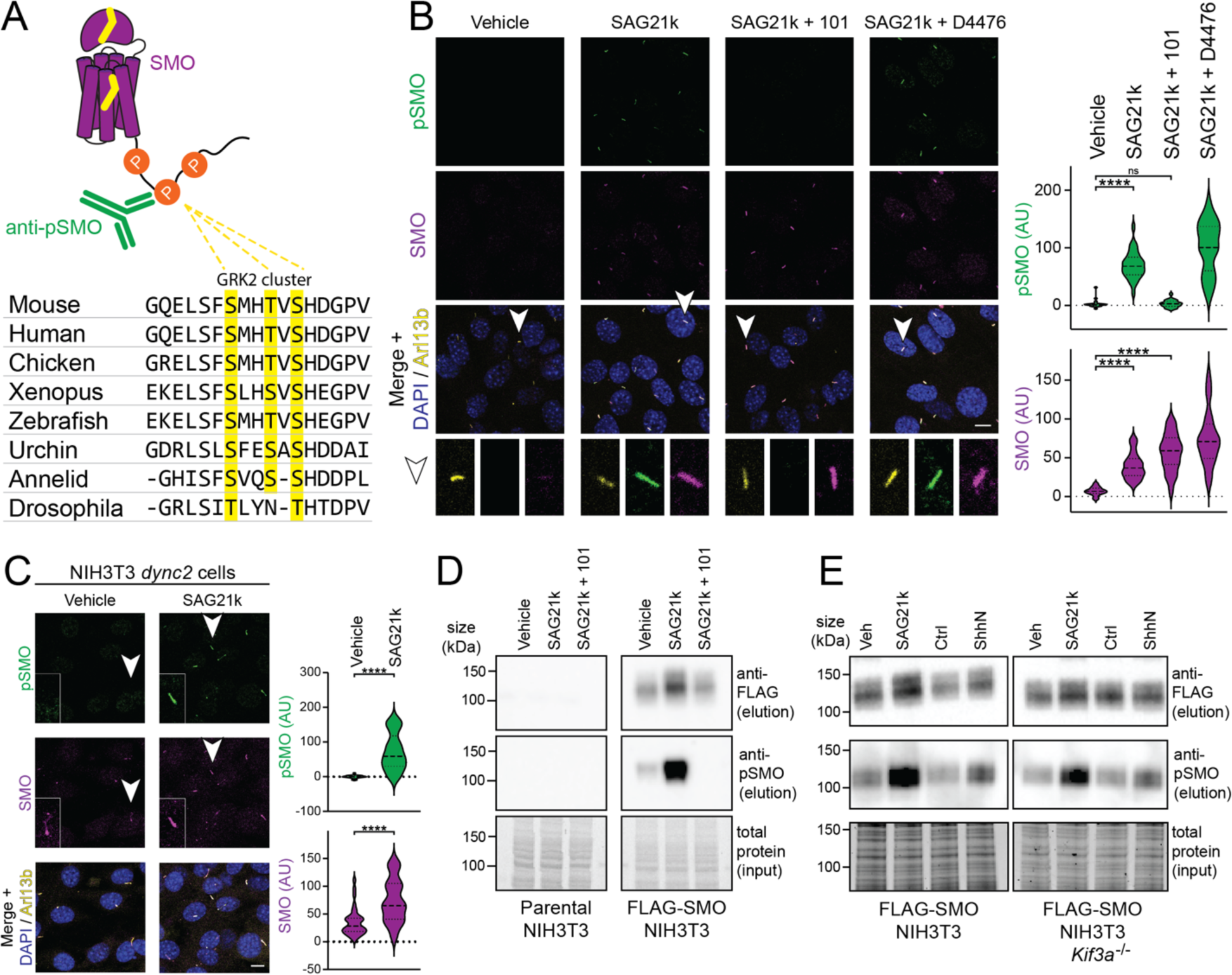
Hh pathway activation leads to accumulation of GRK2-phosphorylated SMO in the cilium. **(A)** Alignment of a region of the SMO pCT from various species, with the yellow highlighted residues indicating the conserved GRK2 phosphorylation cluster recognized by our anti-pSMO antibody (see “Methods”). **(B)** NIH3T3 cells were treated overnight with SAG21k (500 nM) in the presence or absence of the GRK2 inhibitor Cmpd101 (101, 30 µM) or the casein kinase 1 inhibitor D4476 (10 µM), or with a vehicle control. Cells were then fixed and stained with anti-pSMO (green), anti-SMO (magenta), anti-Arl13b (to mark cilia, yellow), and DAPI (to mark nuclei, blue). Representative cilia (arrowhead) are highlighted below each image. **(C)** NIH3T3 *dync2* cells were treated with SAG21k or a vehicle control, then stained and quantified as in (B). Insets show close-up views of a representative cilium from each field. **(D)** NIH3T3 cells stably expressing FLAG-tagged SMO (right) or parental NIH3T3 controls (left) were treated for four hours with vehicle, SAG21k, or SAG21k plus Cmpd101, as described in (B), then lysed and subjected to FLAG immunoaffinity chromatography. FLAG eluates were blotted with anti-pSMO and anti-FLAG to detect phosphorylated SMO and total SMO, respectively. The total protein from the input fractions (prior to FLAG chromatography) serve as loading controls. See Fig. S5H for quantification. **(E)** Isogenic FLAG-SMO-expressing NIH3T3 Flp-in wild-type (left panel) or *Kif3a^-/-^* (right panel) cells^72^ were treated with SAG21k (vs. a vehicle control, lanes 1-2 in each panel), or ShhN conditioned medium (vs. control conditioned medium, lanes 3-4 in each panel), and SMO phosphorylation was monitored via anti-pSMO immunoblotting as in (D). Significance in (B) and (C) was determined via a Mann-Whitney test. *, p <0.05; ****, p < 0.0001; ns, not significant; . n = 22-30 or 32-37 individual cilia per condition in (B) and (C), respectively. Scale bar = 10 μm in all images.

We turned to NIH3T3 fibroblasts to study GRK2 phosphorylation of endogenous SMO in its native ciliary context. These cells are widely considered the “gold standard” cultured cell model for Hh signal transduction, as they are uniformly ciliated and endogenously express all pathway components required to transmit Hh signals from the cell surface to the nucleus^30,31,55,56^. In the Hh pathway “off” state, we did not observe SMO phosphorylation in cilia, as assessed via immunofluorescence microscopy with the anti-pSMO antibody (Fig. 2B). In contrast, endogenous SMO is phosphorylated in cilia following Hh pathway activation using SAG21k (Fig. 2B, Fig. S4D) or the N-terminal signaling fragment of Sonic hedgehog (ShhN) (Fig. S4D). This ciliary SMO phosphorylation was absent in *Smo^-/-^* fibroblasts (Fig. S5A), demonstrating that the anti-pSMO antibody specifically detects endogenous phosphorylated SMO in cilia. The anti-pSMO signal colocalized with the signal from an anti-SMO antibody that stains the ciliary base and shaft regions (and recognizes SMO regardless of phosphorylation state), suggesting that phosphorylated SMO localizes throughout the cilium. Ciliary anti-pSMO staining in NIH3T3 cells was abolished by treatment with each of two chemically distinct GRK2 inhibitors, Cmpd101 or 14as^57^ (Fig. 2B, S5B), or by CRISPR-mediated disruption of *Grk2* (Fig. S5C-E). The effects of GRK2 inhibition were not attributable to changes in SMO ciliary localization, since total SMO in cilia was unchanged by treatment with the GRK2 inhibitors (Fig. 2B, S5B). This finding is consistent with previous observations that GRK2 inhibitors block Hh signal transduction in mouse fibroblasts without affecting levels of SMO in cilia^43–45^. Thus, GRK2 controls SMO phosphorylation during endogenous Hh signal transduction in cilia, at least at the site(s) recognized by our anti-pSMO antibody.

Prior studies have implicated the ɑ and ɣ isoforms of casein kinase 1 (CK1) in SMO phosphorylation^58,59^. These studies initially proposed SMO S594/T597/S599 as CK1 phosphorylation sites, based on an *in vitro* phosphorylation assay using purified CK1ɑ and a soluble construct encompassing the full-length, unstructured SMO cytoplasmic domain^58^. However, compound D4476, which potently inhibits all CK1 isoforms^60–62^, failed to decrease SAG21k-induced phosphorylation of endogenous full-length SMO in NIH3T3 cilia (Fig. 2B), indicating that CK1 does not phosphorylate SMO S594/T597/S599 under physiological conditions.

We considered whether ciliary SMO phosphorylation arises simply from SMO ciliary localization, rather than SMO activation. To distinguish between these possibilities, we uncoupled SMO ciliary trafficking from activation using NIH3T3 cells stably expressing a short hairpin RNA against the dynein 2 heavy chain motor (hereafter referred to as NIH3T3 *dync2* cells)^32,63^, which harbor high levels of inactive SMO in cilia in the pathway “off” state^32,63^ (Fig. 2C). If SMO phosphorylation is a result of SMO localization in cilia, it should occur in NIH3T3 *dync2* cells even without Hh pathway activation. However, we did not detect phosphorylated SMO in vehicle-treated NIH3T3 *dync2* cells, whereas SAG21k treatment strongly induced SMO phosphorylation (Fig. 2C). Furthermore, treatment of wild-type NIH3T3 cells with the SMO inverse agonist cyclopamine, which causes SMO to accumulate in cilia in an inactive conformation^32,64,65^, also failed to induce SMO phosphorylation (Fig. 3B). Thus, SMO must be in an active conformation to undergo GRK2 phosphorylation in the cilium.

**Figure 3:**
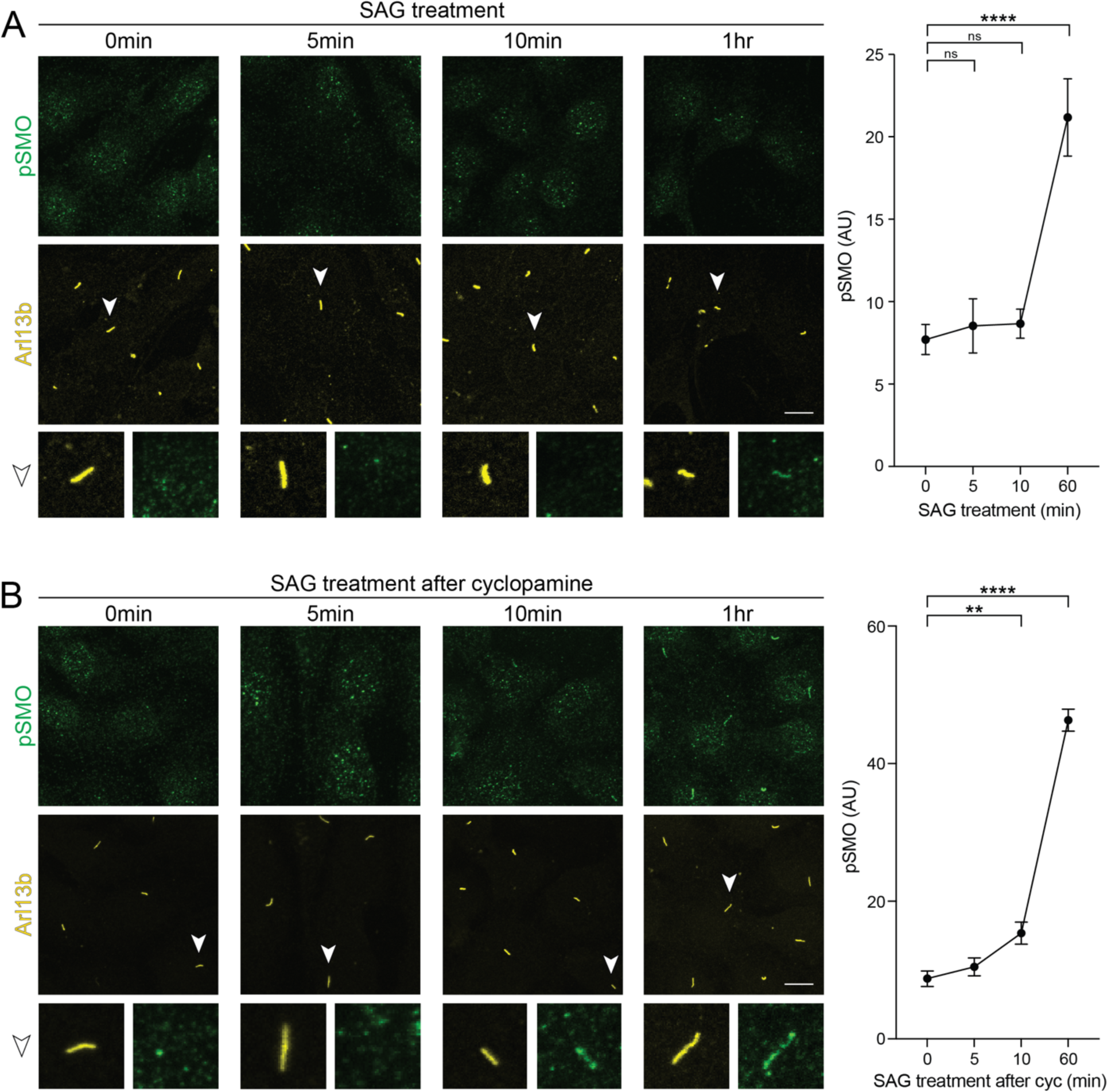
GRK2 phosphorylation of ciliary SMO is an early event in intracellular Hh signal transmission. **(A)** Left: Wild-type NIH3T3 cells were treated with 100 nM SAG for times ranging from 5 minutes to 1 hour, then fixed and stained for pSMO (green) and Arl13b (yellow). Right: quantification of the mean pSMO intensity in cilia at each indicated time point. **(B)** The same experiment was performed, except NIH3T3 cells were first pretreated with 5μM cyclopamine overnight, followed by SAG treatment as described in (A). Representative cilia (arrowhead) are highlighted below each image. n= 50-100 cells per condition. Significance was determined via one-way ANOVA followed by Dunnett’s multiple comparison test. **, p < 0.01; ****, p < 0.0001; ns, not significant. Data are represented as means ± SEM. Scale bar = 5 μm in all images.

To validate the findings from our microscopy studies, we examined SMO phosphorylation in NIH3T3 cells via immunoblotting. Because the anti-pSMO antibody was not sufficiently sensitive to detect phosphorylation of endogenous SMO by immunoblotting (Fig. 2D, “parental”), we generated an NIH3T3 line stably expressing low levels of FLAG-tagged SMO for these studies (Fig. S5F-G), thereby minimizing possible overexpression artifacts. Control experiments confirmed that the stably expressed FLAG-SMO, like its endogenous counterpart, underwent physiological regulation by the ciliary trafficking machinery (Fig. S5G). FLAG-SMO exhibited low but detectable basal phosphorylation that was strongly increased by SAG21k, and this effect was abolished by Cmpd101 (Fig. 2D, “FLAG-SMO”). We note that FLAG-SMO levels increase modestly following SMO activation, consistent with observations regarding endogenous SMO^64^. However, after measuring the normalized ratio of phosphorylated to total SMO in each condition, we conclude that this modest increase in FLAG-SMO levels cannot explain the dramatic effects of SMO agonist or GRK2 inhibitor on SMO phosphorylation that we observed (Fig. 2D, S5H).

To assess whether phosphorylation of active SMO requires an intact primary cilium, we monitored SMO phosphorylation in cells lacking the anterograde kinesin motor KIF3A, a key component of the intraflagellar transport (IFT) machinery needed for ciliary biogenesis and maintenance. Hh signal transduction is disrupted in *Kif3a*^-/-^ cells^66,67^, and these cells fail to form a ciliary shaft, although the ciliary base is comparatively less affected^68–71^. Using immunoblot analysis, we found that either SAG21k or ShhN can still induce SMO phosphorylation in *Kif3a*^-/-^ cells (Fig. 2E). Thus, SMO phosphorylation can occur even when the ciliary shaft is not fully intact, consistent with SMO activation occurring in or near the base of the cilium (see Discussion.)

Together, our immunofluorescence and immunoblotting studies demonstrate that active SMO undergoes GRK2-dependent phosphorylation in ciliated cultured cells.

### GRK2 phosphorylation of ciliary SMO is an early event in Hh signal transmission

While Hh signal transduction pathway steps downstream of SMO, such as post-translational modification of GLI proteins and GLI accumulation at the cilium tip, begin to occur within 1-2 hours of SMO activation^22,24,73^, the kinetics of SMO phosphorylation and SMO / PKA-C interaction in cilia are unknown. We used our anti-pSMO antibody to assess whether SMO undergoes phosphorylation in cilia on a time scale consistent with established Hh pathway signaling events.

In initial experiments, we observed SMO phosphorylation in NIH3T3 cilia starting one hour after pathway activation (Fig. 3A). However, these measurements likely underestimate the speed of ciliary SMO phosphorylation because SMO is present at extremely low levels in cilia during the initial phases of Hh pathway activation and requires several hours to accumulate to high levels in this compartment^31,32^. Our experiments might therefore fail to capture SMO phosphorylation at early time points when SMO levels are below the threshold for reliable detection.

To circumvent this issue, we used two independent strategies to enable detection of SMO phosphorylation at early stages of Hh pathway activation, and thereby unmask the true kinetics of SMO phosphorylation in cilia. Critically, both approaches measure endogenous SMO protein, thereby avoiding potential pitfalls of overexpression-based strategies. First, we treated NIH3T3 cells with the SMO inverse agonist cyclopamine to trap SMO in an inactive, nonphosphorylated state in cilia^32,39,64,65^, then rapidly activated SMO by replacing cyclopamine with the SMO agonist SAG. Using this approach, we detected SMO phosphorylation in cilia after only 10 minutes of SAG treatment (Fig. 3B. S6A). Second, in NIH3T3 *dync2* cells, which display some degree of SMO ciliary localization even in the pathway “off” state^32,63^ (Fig. 2C), we detected SMO phosphorylation in cilia within 5 minutes of SAG21k treatment (Fig. S6B). Thus, ciliary SMO undergoes phosphorylation within minutes of Hh pathway activation, consistent with a role for SMO phosphorylation in initiating the intracellular steps of Hh signal transduction.

### SMO phosphorylation in the ciliary membrane requires continuous GRK2 activity but not heterotrimeric Gýy subunits

Once SMO is activated and undergoes GRK2-mediated phosphorylation, it may persist in a phosphorylated state within the cilium for many hours, without requiring additional GRK2 activity for maintenance. Alternatively, phosphorylation may be dynamic and labile, requiring continuous GRK2 kinase activity to maintain the pool of ciliary SMO in its phosphorylated state. To distinguish between these possibilities, we utilized Cmpd101 to determine how blockade of GRK2 kinase activity impacts the amount of phosphorylated SMO. In immunofluorescence microscopy, SAG21k-induced phosphorylation of ciliary SMO decreased rapidly following Cmpd101 treatment, reaching half-maximal intensity after 15 minutes and returning to baseline levels by 2 hours (Fig. 4A). Cmpd101 induced an even faster disappearance of phosphorylated SMO when measured by immunoblotting, reaching half-maximal intensity after 5 minutes and undetectable levels at 1 hour (Fig. 4B). These studies indicate that continuous GRK2 kinase activity maintains a pool of ciliary SMO in a phosphorylated state.

**Figure 4:**
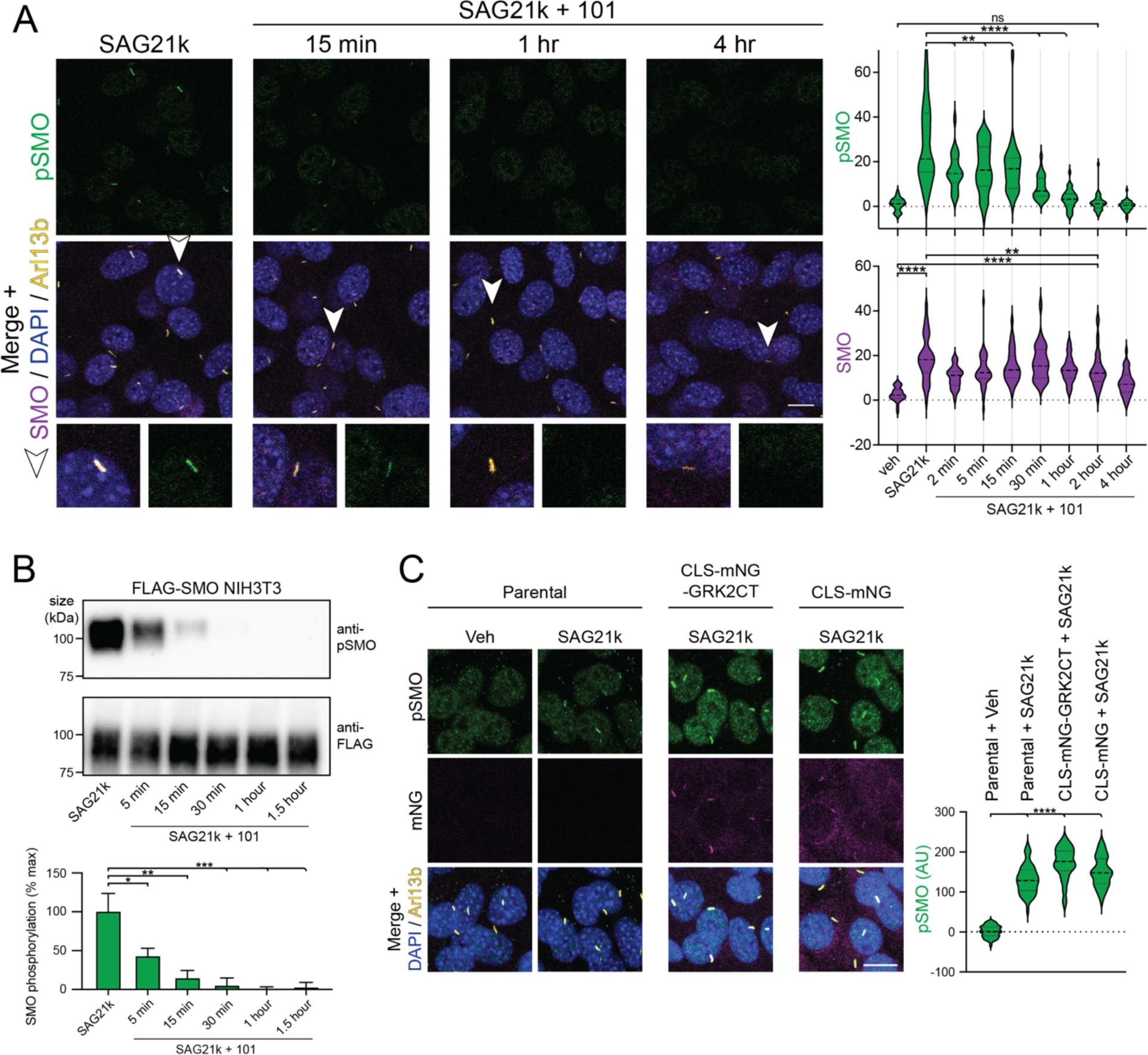
Continuous GRK2 activity sets the SMO phosphorylation state. **(A)** NIH3T3 cells were treated overnight with SAG21k (500 nM) (left-most panel), and then Cmpd101 (101, 30 µM) was added for the indicated times. Cells were then fixed and stained for pSMO (green), SMO (magenta), Arl13b (yellow), and DAPI (blue), as described in Fig. 2B. Representative cilia (arrowhead) are highlighted below each image. Quantification of pSMO and total SMO at the indicated time points are shown below. **(B)** NIH3T3 cells stably expressing FLAG-tagged SMO were treated as in (A), then SMO was purified using FLAG affinity chromatography and the eluates blotted with anti-pSMO and anti-FLAG antibodies. Graph below indicates quantification of band intensities. **(C)** A construct consisting of GRK2CT fused to a ciliary targeting signal and mNeonGreen (mNG) fluorescent protein^52^, or a control construct lacking GRK2CT, was stably expressed in NIH3T3 Flp-in cells. Following SAG21k treatment, SMO phosphorylation (green), Arl13b (yellow), DAPI (blue), and the cilium-targeted fusion (mNG, magenta) in the indicated stable cell lines (middle and right panels), or parental NIH3T3 Flp-in control cells (left panels), were monitored via immunofluorescence. Significance in (A) and (C) was determined via a Mann-Whitney test, and in (B) via one-way ANOVA. *, p < 0.05; **, p < 0.01; ****, p < 0.0001; ns, not significant; n = 24-47 individual cilia per condition in (A); mean +/-standard deviation is shown in (B). Scale bar = 10 μm.

We next investigated how GRK2 recognizes activated SMO in the cilium, specifically, whether its recruitment mechanism mirrored that of classical GPCRs in the plasma membrane. The ý and ψ subunits of heterotrimeric G proteins (Gýy) recruit GRK2 to the plasma membrane via direct interactions with the GRK2 C-terminus (GRK2CT)^74,75^. Accordingly, expression of GRK2CT strongly reduces GRK2 phosphorylation of plasma membrane GPCRs, via dominant-negative blockade of Gýy interactions with endogenous GRK2^74–76^. In contrast, we observed no impact on SMO phosphorylation in the cilium when we targeted GRK2CT to this compartment (Fig. 4C, see Methods). These data demonstrate that Gýy subunits are not required for GRK2 to recognize and phosphorylate activated SMO in cilia.

### GRK2 phosphorylation controls SMO / PKA-C colocalization in cilia

Given that SMO undergoes activity-dependent GRK2 phosphorylation in cilia, we asked whether GRK2 phosphorylation precedes and is required for interaction with PKA-C in this organelle. The amount of PKA-C in cilia is vanishingly small, based on microscopy and proximity-based mass spectrometry measurements^39,51,77,78^. To facilitate PKA-C detection in cilia, we therefore increased the amount of its key interacting partner, SMO, in the ciliary membrane by utilizing a stable NIH3T3 cell line expressing V5-TurboID-tagged SMO at low levels; the additional ciliary SMO protein helps to “kinetically trap” PKA-C in this compartment (see Discussion), raising SMO / PKA-C levels above the limit of detection in microscopy^39^. We previously used this strategy (and this cell line) to demonstrate that SAG-bound SMO colocalizes with PKA-C in cilia but cyclopamine-bound SMO does not, confirming that SMO / PKA-C colocalization depends strictly on SMO activity state and ruling out potential nonspecific effects of SMO overexpression on SMO/ PKA-C ciliary colocalization^39^.

SMO / PKA-C colocalization occurred with somewhat slower kinetics than SMO phosphorylation but was nevertheless readily detectable after one hour of SAG treatment (Fig. 5A), consistent with SMO phosphorylation preceding PKA-C recruitment during this process. Critically, SAG-induced SMO / PKA-C colocalization in cilia was abolished by treatment with either Cmpd101 or 14as (Fig. 5B, S7). These studies demonstrate that GRK2 phosphorylation controls SMO / PKA-C colocalization in cilia.

**Figure 5:**
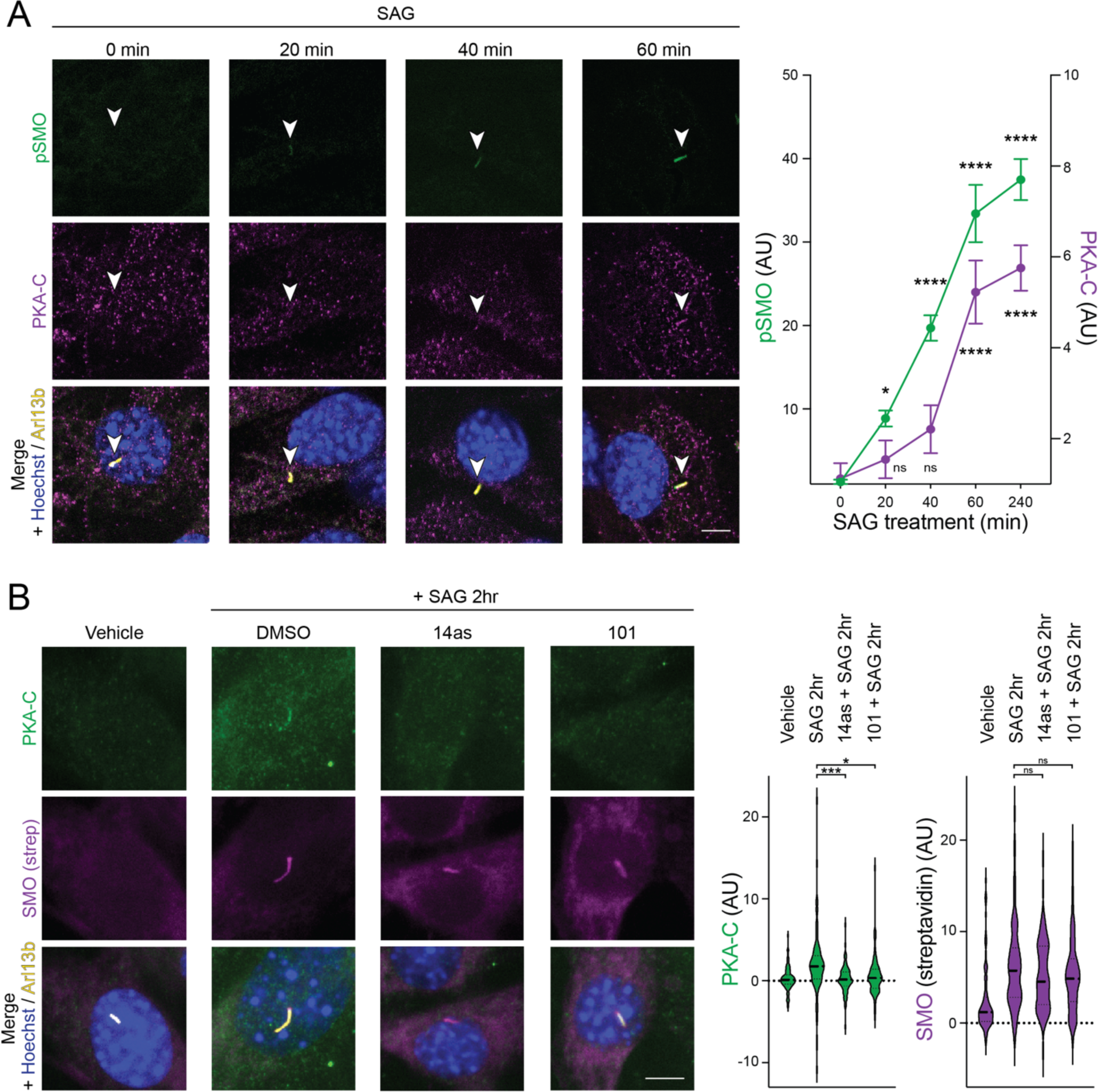
GRK2 phosphorylation controls SMO / PKA-C colocalization in cilia. **(A)** NIH3T3 cells stably expressing SMO-V5-TurboID were pretreated with cyclopamine for 16hr, followed by SAG for the indicated times, and accumulation of PKA-C (magenta) and phosphorylated SMO (pSMO, green) in cilia (Arl13b, yellow) were monitored via immunofluorescence microscopy. Hoechst (blue) marks nuclei. Left: raw data; Right: quantification of pSMO (left axis) and PKA-C (right axis). Cilium is marked by an arrowhead. **(B)** NIH3T3 cells stably expressing SMO-V5-TurboID were treated with vehicle, Cmpd101 or 14as for 16 hours. SAG was then added to the culture medium in the presence of the indicated inhibitors and incubated for 2 hours. Cells were labeled with biotin for 10 min before fixation, and stained for PKA-C (green), streptavidin (magenta), Arl13b (yellow), and Hoechst (blue). Streptavidin staining marks the localization of SMO-V5-TurboID in the cilium. Left: raw data; Right: quantification. Significance was determined via one-way ANOVA followed by Dunnett’s multiple comparison test. **, p < 0.01; ****, p < 0.0001; ns, not significant; n = 50-100 cells per condition. Data are represented as means ± SEM. Scale bar = 5 μm in all images.

### GRK2 phosphorylation of SMO is sufficient to induce SMO / PKA-C interaction

We next sought to uncover the biochemical mechanism by which GRK2 phosphorylation promotes formation of SMO / PKA-C complexes. We considered two possible models. Phosphorylation of SMO may trigger a direct interaction with PKA-C (Fig. 6A, “Direct Model”). Alternatively, phosphorylation may recruit additional proteins that promote the SMO / PKA-C interaction (Fig. 6A, “Indirect Model”).

**Figure 6:**
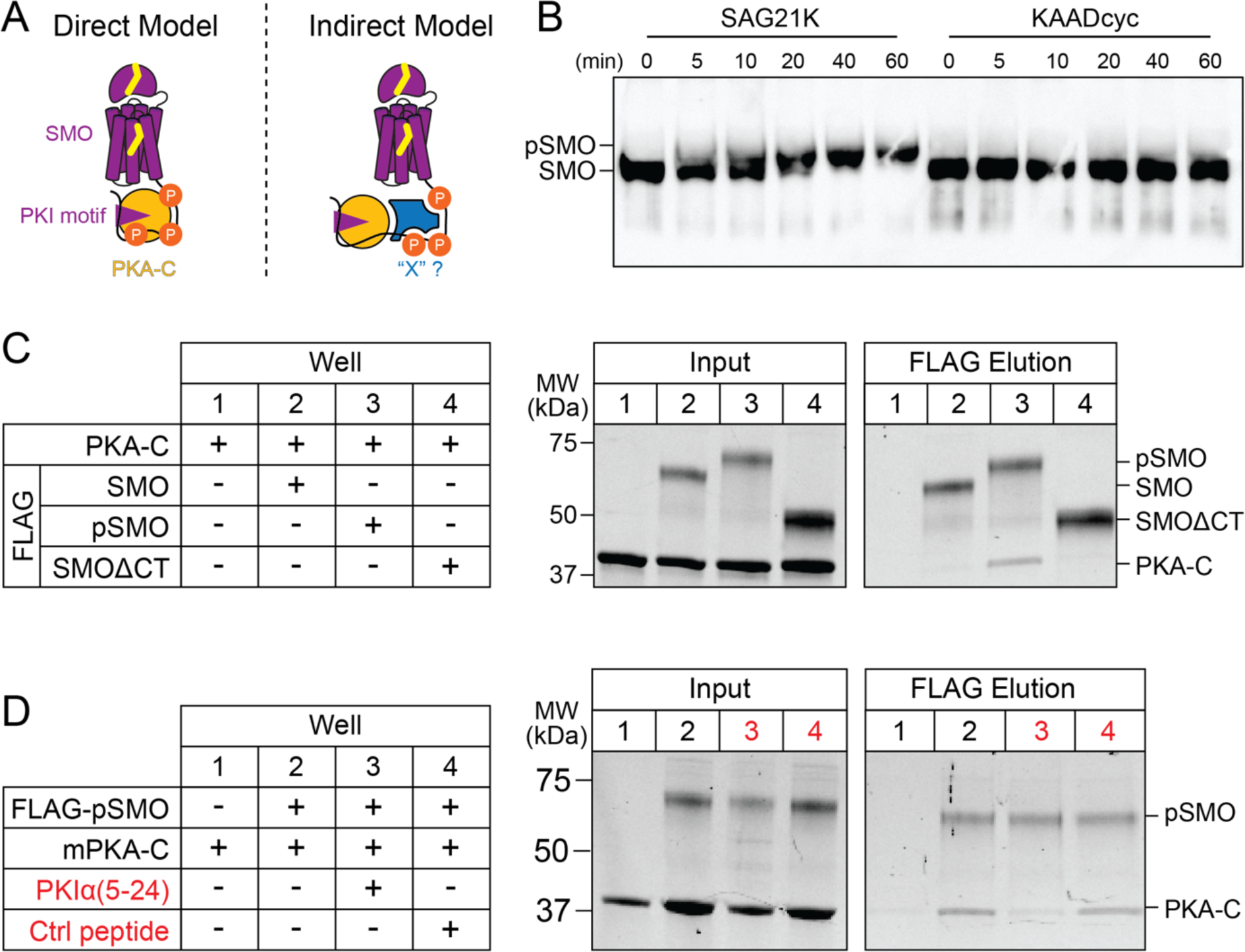
GRK2 phosphorylation of SMO is sufficient to induce SMO / PKA-C interaction. **(A)** Schematic diagram for two models of how GRK2 phosphorylation of SMO may promote interactions with PKA-C, either by directly triggering formation of a SMO / PKA-C complex (Direct Model) or acting via an intermediary protein, symbolized as “X” (Indirect Model). **(B)** Purified FLAG-tagged SMO was subject to phosphorylation by GRK2 *in vitro* in the presence of SMO agonist (SAG21k) or inverse agonist (KAADcyc) for the indicated minutes (min), and conversion of SMO to a phosphorylated form (pSMO) was analyzed by monitoring its mobility shift on SDS-PAGE via anti-FLAG immunoblotting. **(C)** Purified FLAG-tagged SMO in either a nonphosphorylated or phosphorylated state (SMO or pSMO, respectively, at 18.2 µM), was mixed with PKA-C *in vitro* (10 µM), and complex formation was detected via pulldown on anti-FLAG beads, followed by analysis of total protein in input and FLAG elution fractions on SDS-PAGE via Stain-free imaging. SMO lacking the cytoplasmic tail (FLAG-SMOΔCT) serves as a negative control for PKA-C binding. Positions of SMO, pSMO, SMOΔCT, and PKA-C are indicated at right. **(D)** SMO / PKA-C pulldowns were performed as in (C), except that the following competitor peptides (red) were added during the initial SMO / PKA-C binding step: either a PKA pseudosubstrate peptide from PKIα (PKIα(5-24)), or a control, non-PKA-binding peptide (Ctrl peptide), each at a final concentration of 100 μM.

To distinguish between these scenarios, we biochemically reconstituted SMO / PKA-C complexes using purified proteins *in vitro*. Biochemical reconstitution is uniquely suited to address these models because it enables a stringent definition of the minimal set of factors required for GRK2 phosphorylation to mediate SMO / PKA-C interactions. In contrast, such questions are nearly impossible to address in the complex, biochemically undefined environments of cells or organisms, where proteins, lipids, metabolites, and other molecules or ions may contribute to interactions between GRK2-phosphorylated SMO and PKA-C.

We prepared purified SMO protein in GRK2-phosphorylated or non-phosphorylated states, and then evaluated the ability of each of these preparations to bind purified PKA-C *in vitro.* To obtain GRK2-phosphorylated SMO, we established procedures to purify near-full-length FLAG-tagged SMO including the extracellular cysteine rich domain (CRD), 7TM domain, and pCT (Fig. S8A), then optimized conditions to phosphorylate SMO *in vitro* using purified GRK2 (see Methods). Following incubation with GRK2, we detected a strongly phosphorylated SMO species in the presence of SMO agonist but not SMO inverse agonist, manifest as incorporation of γ^32^P-ATP in autoradiography (Fig. S8B) and decreased electrophoretic mobility on SDS-PAGE (Fig. 6B). These results are consistent with GRK2’s preference for the SMO active conformation under physiological conditions^38,46^ (Fig. 2). We confirmed via anti-pSMO immunoblotting that SMO underwent phosphorylation *in vitro* at the physiological GRK2 cluster (Fig. S8C). To prepare non-phosphorylated SMO, we expressed and purified SMO in the presence of an inverse agonist, thereby avoiding phosphorylation by endogenous GRK2 in our HEK293 expression system, and omitted the *in vitro* GRK2 phosphorylation step.

To evaluate whether SMO phosphorylation is sufficient to enable binding to PKA-C, we mixed phosphorylated or nonphosphorylated SMO (Fig. S8D) with PKA-C in a buffer containing glyco-diosgenin (GDN) detergent to keep SMO soluble, SAG21k to maintain SMO in an active conformation, and Mg/ATP to promote PKA-C pseudosubstrate interactions^39^, followed by FLAG affinity chromatography. Only trace amounts of PKA-C were bound by non-phosphorylated SMO, similar to those pulled down by a negative control SMO construct lacking the entire cytoplasmic domain (SMOΔCT, Fig. 6C). This result is consistent with the relatively modest affinity of PKA-C for a soluble, nonphosphorylated SMO pCT construct in surface plasmon resonance studies (KD = 752 +/-34 nM)^39^. In contrast, phosphorylated SMO readily bound PKA-C (Fig. 6C). Thus, GRK2 phosphorylation of SMO can enhance SMO / PKA-C interactions with no other proteins present. Binding between phosphorylated SMO and PKA-C required the SMO PKI motif, as it was reduced to near-background levels by a peptide from the PKA-C pseudosubstrate PKIα that blocks the PKA-C active site^79^ and competes with the SMO PKI motif^39^ (Fig. 6D). We note that SMO binds substoichiometric amounts of PKA-C in this assay (FLAG elutions contain 25 +/-6.8% as much PKA-C as SMO, n = 4 independent experiments), suggesting that additional factors present in living systems may increase the efficiency of the interactions we observe *in vitro* (see Discussion). Nevertheless, our *in vitro* experiments indicate that such factors are not absolutely required, leading us to conclude that GRK2 phosphorylation alone is sufficient for direct SMO / PKA-C interactions (Fig. 6A, “direct”).

### GRK2 phosphorylation of ciliary SMO is a general, evolutionarily conserved component of Hh signal transduction *in vivo*

We have shown that the ciliary SMO-GRK2-PKA communication pathway operates in fibroblast cell lines, but whether this pathway operates during Hh signal transduction in other biological contexts is unknown. To study GRK2 phosphorylation of ciliary SMO during Hh signaling *in vivo,* we utilized our anti-pSMO antibody to ask whether SMO undergoes GRK2-mediated phosphorylation in a range of cellular and *in vivo* contexts. We focused primarily on the nervous system, where Hh signaling plays numerous well-established roles in proliferation and differentiation, both during embryogenesis and postnatally^80–82^.

We first utilized the vertebrate embryonic neural tube to study the instructive role of Hh signaling in cell fate decisions during neural development. This structure, the precursor to the spinal cord, is patterned by Sonic hedgehog (Shh) produced by the notochord and floor plate, which diffuses along the dorsal-ventral axis to form a gradient that instructs neuronal cell fate decisions in a concentration-and duration-dependent fashion^83,84^. When applied to neural tube sections from wild-type mice, the anti-pSMO antibody prominently stained cilia in the ventral neural tube, where levels of Shh are highest (Fig. 7A), but not in the dorsal neural tube where levels of Shh are low (Fig. S9A). The anti-pSMO stain observed in wild-type mice (*Smo^WT^*) was a faithful measure of SMO activation, as the signal was noticeably weaker in a *Smo* hypomorphic mutant (*Smo^cbb^*), considerably stronger in a *Ptch1* null mutant (*Ptch1^-/-^)* or a constitutively activating *Smo* gain-of-function mutant (*Smo^M^*^2^), and absent in a *Smo* null mutant (*Smo^bnb^*)^85–88^ (Fig. 7A). To determine whether SMO is phosphorylated during neural development in other vertebrate species, we stained whole-mount zebrafish embryos (24 hours post-fertilization) with our anti-pSMO antibody. Again, we observed an intense anti-pSMO staining in cilia at the ventral but not the dorsal spinal cord, representing a population of neuronal progenitors that require high levels of Shh to generate primary motor neurons (pMN) and oligodendrocytes^89–91^ (Fig. 7B). Thus, SMO undergoes activity-dependent GRK2 phosphorylation during instructive Hh signaling in vertebrate neural tube development.

**Figure 7:**
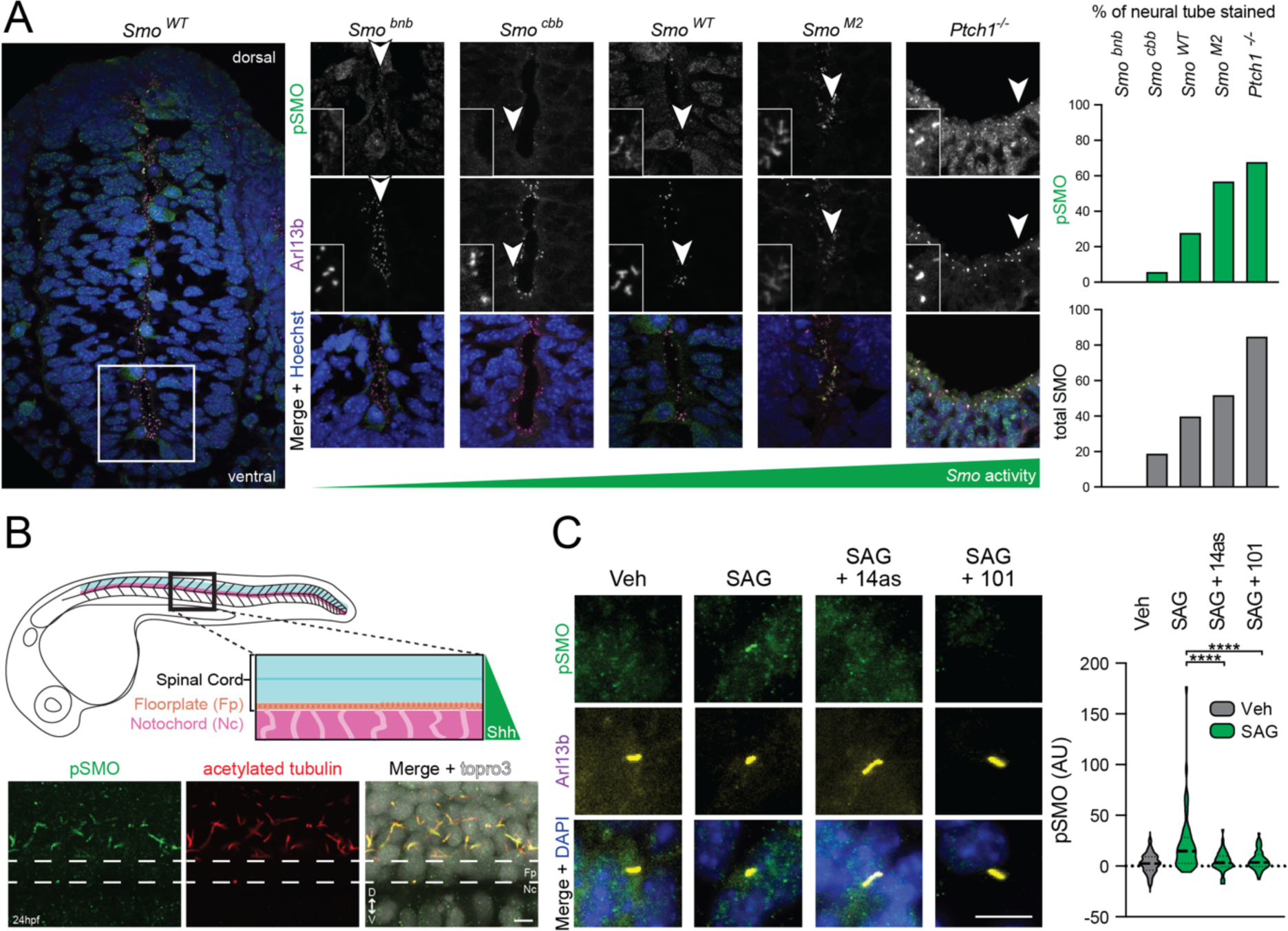
GRK2 phosphorylation of ciliary SMO is a general, evolutionarily conserved aspect of Hh signal transduction *in vivo*. **(A)** Mouse neural tubes from wild-type, *Smo,* or *Ptch1* mutant E9.5 embryos (low-magnification view of wild-type mouse at left, higher-magnification view of wild-type, *Ptch1^-/-^,* or *Smo* alleles of differing strengths at right) were stained for pSMO (green), cilia (Arl13b, magenta), and nuclei (Hoechst, blue). Images are oriented with dorsal pointing up and ventral pointing down. Insets show close-up views of representative cilia from each field. The percent of each neural tube that was stained by anti-pSMO (green) or anti-SMO (gray) antibodies is quantified at right. Raw data from anti-SMO staining is presented in Fig. S9B. **(B)** Top: Schematic of 24 hpf zebrafish embryos showing location of the spinal cord relative to Shh-secreting floorplate and notochord. Bottom: Lateral views of whole-mount zebrafish embryos at 24 hours post-fertilization (hpf) were stained with anti-pSMO (green) or anti-acetylated tubulin (red) antibodies, with topro3 counterstain to mark nuclei (white). Magnified view of the ventral spinal cord is shown, with the floorplate indicated by dashed white lines. The dorsal spinal cord is facing up and the ventral spinal cord is facing down. **(C)** CGNPs freshly isolated from neonatal mice were cultured *ex vivo* and treated with vehicle or SAG, in the presence or absence of Cmpd101 or 14as, followed by staining with anti-pSMO (green) and Arl13b (yellow). Ciliary anti-pSMO intensity is quantified and significance determined via one-way ANOVA. ****, p < 0.0001. Scale bar = 5 μm in (B); 10 μm in (C).

We extended our analysis of SMO phosphorylation to the early zebrafish brain and eye field primordium, both of which are patterned by Shh produced at the embryonic midline^92–94^. Here we observed sparse but detectable ciliary anti-pSMO staining that was dramatically enhanced by treatment of embryos with SAG21k, demonstrating endogenous SMO phosphorylation in these structures and consistent with the ability of this SMO agonist to ectopically activate Hh signaling during embryogenesis (Fig. S10)^95–99^. Thus, SMO undergoes GRK2 phosphorylation during eye and brain development.

Lastly, we turned to the proliferation of mouse cerebellar granule neural precursors (CGNP) in culture as a model for mitogenic Hh signaling in the postnatal nervous system. Shortly after birth, Shh produced by the Purkinje cell layer of the cerebellum serves as the major mitogen for the adjacent CGNPs, driving Hh signal transduction that leads to transcription of genes involved in cell cycle entry and ultimately fueling a nearly one thousand-fold expansion of the postnatal cerebellum^100–103^. This phenomenon can be recapitulated in culture, where CGNPs dissected from postnatal mice and cultured *ex vivo* mount a robust proliferative response to Hh pathway stimulation that requires SMO activity and primary cilia. Primary CGNP cultures treated with SAG but not a vehicle control displayed strong ciliary anti-pSMO staining that was blocked by Cmpd101 or 14as, demonstrating that GRK2 phosphorylates active SMO during CGNP proliferation (Fig. 7C, S11).

Taken together, our findings in mouse and zebrafish neural progenitors, zebrafish eye and forebrain precursor cells, and mouse CGNPs demonstrate that GRK2 phosphorylation of SMO is a general, evolutionarily conserved aspect of vertebrate Hh signal transduction.

## DISCUSSION

SMO inhibition of PKA-C is fundamental to Hh signal transduction in development, homeostasis, and disease^2–4,34^. Here we identify GRK2 as an indispensable regulator of this inhibition during Hh signal transduction in primary cilia. We show that upon Hh pathway activation, ciliary GRK2 rapidly recognizes the active, agonist-bound conformation of SMO, and subsequently phosphorylates essential sites in the SMO pCT. This phosphorylation is: 1) sufficient to trigger direct SMO / PKA-C interactions, 2) required for SMO to engage PKA-C in cilia, and 3) a general feature of SMO activation by naturally occurring and synthetic SMO agonists, and in multiple cellular and *in vivo* contexts. Together with prior functional studies^38,39,43–45^, our work establishes GRK2 phosphorylation of SMO, and the ensuing PKA-C recruitment and inactivation, as critical initiating events for the intracellular steps in Hh signal transduction. These findings provide a deeper mechanistic understanding of canonical Hh signal transduction throughout development and disease. Furthermore, we anticipate that the tools and concepts established here will enable future cell biological investigations of SMO-GLI communication in many tissue, organ, and animal contexts.

### A model for SMO-GRK2-PKA signaling in primary cilia

Based on our findings, we propose the following model for SMO-GRK2-PKA communication during Hh signal transduction (Fig. 8). In the Hh pathway “off” state, GRK2 is poised at the base of the cilium, but SMO is in an inactive conformation and cannot be recognized or phosphorylated by GRK2. As a result, PKA-C remains active as it transits through the cilium and can phosphorylate and thereby trigger inactivation of GLI^20,33,77,78^ (Fig. 8, left panel). In the Hh pathway “on” state, SMO binds sterols and assumes an active conformation; GRK2 recognizes active SMO, resulting in accumulation of GRK2 in the ciliary shaft and phosphorylation of the SMO pCT (Fig. 8, middle panel). Consequently, SMO can bind and inhibit PKA-C in the ciliary shaft, leading to GLI activation (Fig. 8, right panel). These processes are further strengthened by the accumulation of SMO protein to high levels in the cilium over the course of several hours following SMO activation^31,32^, which undergoes GRK2 phosphorylation and subsequently binds and inhibits additional PKA-C molecules transiting through the cilium, thereby serving as a positive feedback loop for the SMO-GRK2-PKA communication pathway described above. Ciliary SMO thereby acts as a “kinetic trap” for PKA-C, causing PKA-C activity in cilia to decrease^104^, even though PKA-C levels in cilia increase (Fig. 5, see also ^39^).

**Figure 8:**
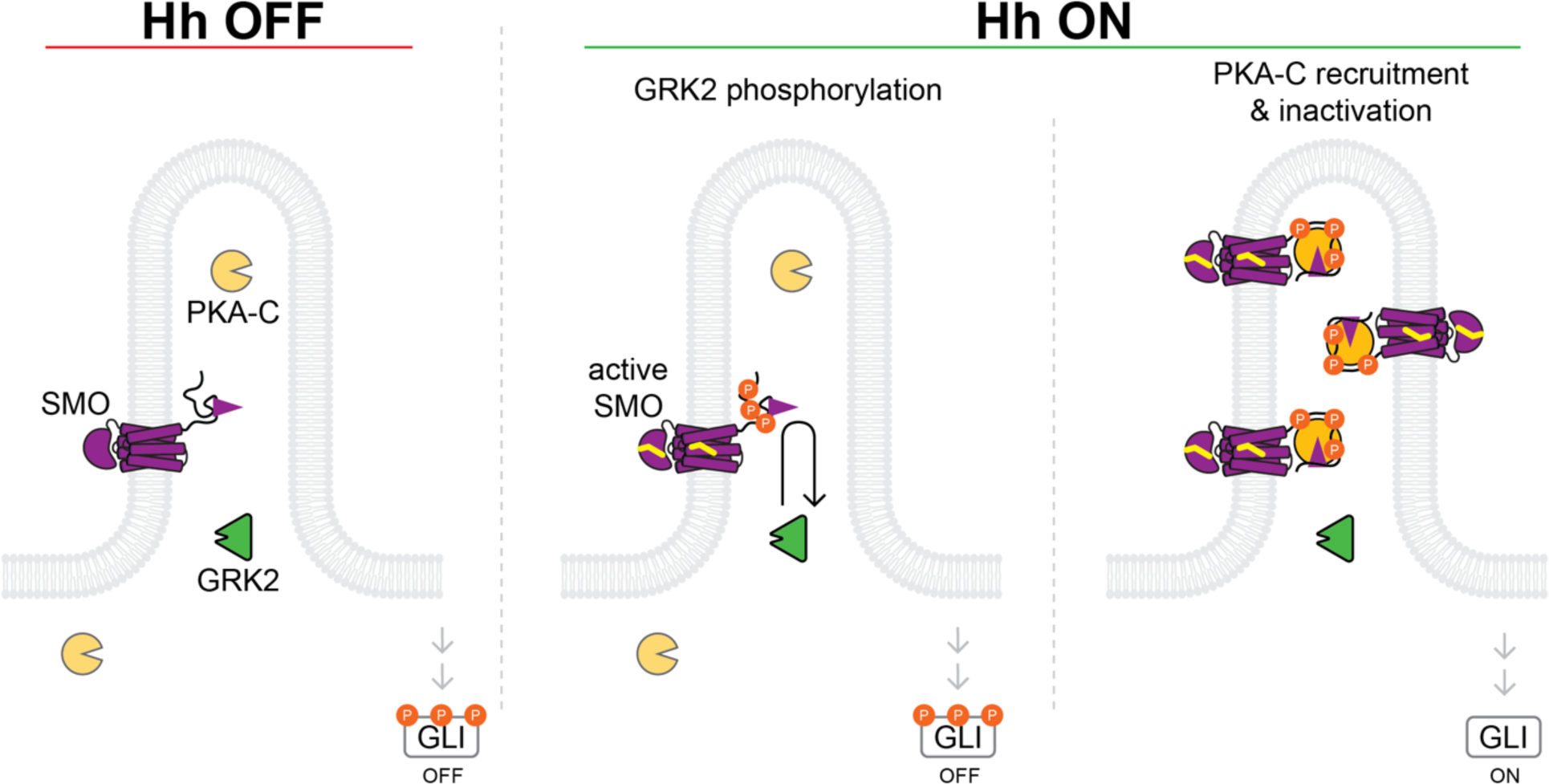
Model for SMO-GRK2-PKA communication during Hh signal transduction in the cilium. See main text for details.

Our prior experiments in cultured fibroblasts and zebrafish embryos support an essential role for the SMO-GRK2-PKA communication pathway in Hh signal transduction^38,39^, but SMO can also affect ciliary PKA via at least two additional mechanisms: 1) coupling to inhibitory G proteins (Gα_i_)^105–107^, which block PKA-C by inhibiting the generaton of cAMP; 2) stimulating the ciliary exit of GPR161, a constitutively active GPCR that recruits PKA holoenzymes to cilia via a C-terminal A-kinase anchoring protein domain, and activates PKA-C by coupling to stimulatory (Gα_s_) G proteins^44,108–112^. These processes, while not absolutely required for SMO-GLI communication^104,113,114^, likely make important, context-dependent contributions to GLI activation, as evidenced by the hyperactive Hh signaling observed in *Gpr161^-/-^*mice^44,108–110^. In sum, the SMO-GRK2-PKA regulatory mechanism, operating in concert with other SMO-PKA signaling pathways, ensures that only the active state of SMO can bind and inhibit PKA-C in the cilium, and thereby helps to avoid pathologic outcomes associated with too much or too little GLI activation^18^.

Our work helps to resolve outstanding questions regarding whether SMO is a target of GRK2 under physiological conditions^43,44,58,115^. Our previous studies implicating GRK2 in SMO / PKA-C signaling relied on reductionist experimental systems that do not fully reflect Hh signal transduction in its native ciliary context. Thus, while SMO undergoes activity-dependent phosphorylation by GRK2 in transfected HEK293 cells^38^, whether this occurs during endogenous Hh signaling in primary cilia was not known. In the present study, we addressed these issues by developing microscopy approaches to sensitively track GRK2 localization over time, as well as an anti-pSMO antibody to directly monitor GRK2 phosphorylation of endogenous SMO in cilia. Using these approaches, we find that GRK2 recognizes and phosphorylates the active conformation of SMO in cilia in several cultured cell and animal systems (paving the way for future studies of GRK2-dependent SMO phosphorylation *in vivo* using genetics or pharmacology). We also show that GRK2 activity is necessary for SMO to engage PKA-C in cilia, echoing the requirement for GRK2 during SMO-GLI communication in cell, organ, and animal models^43–45^. Thus, the current study demonstrates that the SMO-GRK2-PKA signaling pathway underlies physiological Hh signal transduction in primary cilia.

Interestingly, SMO phosphorylation in response to exogenous (SAG21k) or endogenous (ShhN) pathway activators can occur even in *Kif3a*^-/-^ cells, which lack the entire ciliary shaft but maintain at least some aspects of the ciliary base^68–71^. These findings suggest that neither SMO activation nor the ensuing GRK2 phosphorylation requires a fully intact primary cilium. Instead, the ciliary base (where PTCH1 is enriched^31^) may orchestrate these processes, and the ciliary shaft may control Hh pathway steps downstream of SMO. However, two caveats are worth noting. First, our immunoblot experiments in wild-type and *Kif3a*^-/-^ cells necessitated SMO overexpression, and the conclusions may or may not apply to endogenous SMO. Second, we do not yet understand whether components of the ciliary base, or even the cytoplasmic membrane/microtubule network, compensate for the ciliary defects that occur when key ciliary components such as KIF3A are disrupted; as such, the cilium may not be absolutely required for SMO phosphorylation, but it may still serve as the primary location for phosphorylation under physiological conditions. In addition, an alternative model - is that SMO phosphorylation occurs not at the ciliary base, but rather at the plasma membrane, followed by rapid entry of phosphorylated SMO into the primary cilium,although the rapid kinetics of SMO activation-induced ciliary GRK2 recruitment and SMO phosphorylation highlight the cilium as a likely subcellular location for these events. Future studies of SMO phosphorylation in cells lacking key components of the cilium, including the base, shaft, and machinery that enables docking of the mother centriole at the plasma membrane, may help to define the subcellular location of SMO phosphorylation more precisely.

If the ciliary shaft is not involved in GRK2-mediated phosphorylation of active SMO, then why is this structure required for Hh signal transduction, as evidenced by the profound loss of GLI activation in *Kif3a^-/-^* cells and embryos^66,67^? Perhaps the ciliary shaft mediates essential Hh signal transduction events downstream of SMO activation or GRK2-mediated phosphorylation. This may include 1) the interaction between phosphorylated SMO and the pool of active PKA-C that phosphorylates and inhibits GLI; consistent with this possibility, targeted inactivation of PKA-C in the ciliary shaft leads to GLI activation, whereas PKA-C inactivation in other subcellular locations does not^77,78^; 2) addition of activating GLI post-translational modification(s), or removal of inhibitory ones^24,116–118^. Addressing the role of the cilium shaft in these and other regulatory mechanisms represents an important next step for the field.

### Biochemical mechanism of SMO phosphorylation and SMO / PKA-C interaction

Our study also provides new insights into the biochemical mechanism by which GRK2 phosphorylation promotes SMO / PKA-C interactions. We previously showed that GRK2 phosphorylation is necessary for SMO to bind PKA-C in cells^38^, but it was not clear whether phosphorylation directly mediates SMO / PKA-C binding or whether the effect requires additional as-yet-unidentified proteins. We now establish using biochemical reconstitution that GRK2 phosphorylation can enhance binding of purified, near-full-length SMO to purified PKA-C *in vitro*. This result reveals that GRK2 phosphorylation is not merely a permissive event that facilitates the action of another protein on the SMO / PKA-C complex, but is sufficient on its own to trigger SMO / PKA-C binding. In contrast, if the effect of phosphorylation on the interaction needed additional factors, then phosphorylated SMO would not engage PKA-C more efficiently than would non-phosphorylated SMO in our experiments. We note, however, that proteins, lipids, or other factors not present in our *in vitro* system, while not absolutely required, may render phosphorylation-induced SMO / PKA-C binding more efficient in living systems. We expect such factors would play important roles in Hh signaling, and our *in vitro* reconstitution system may enable their discovery and characterization.An outstanding challenge is to understand in structural terms how GRK2 recognizes and phosphorylates the SMO active conformation. Some clues may be provided by recent cryoEM structures of GRK1 in complex with the GPCR rhodopsin. GRK1 recognizes the active conformation of rhodopsin by inserting its N-terminal ɑ-helix (ɑN) into an intracellular rhodopsin cavity exposed by outward movement of TM helices 5 and 6 during rhodopsin activation. This docking event activates GRK1, enabling phosphorylation of rhodopsin intracellular domains, followed by dissociation of the rhodopsin-GRK1 complex^42,119^. We speculate that a similar process underlies GRK2 recognition and phosphorylation of active SMO, as SMO activation causes an outward shift in TM helices 5 and 6^27^, and GRK2 constructs harboring point mutations in the ɑN helix fail to rescue the loss of SMO-GLI communication observed in *Grk2^-/-^* fibroblasts^44^. Nevertheless, other regions of GRK2 critical for GPCR regulation, such as the phosphoinositide- and Gβγ-binding pleckstrin homology domain^40,41^, are not conserved in GRK1^42,119^, and the structural basis for these regions to enable GRK2 phosphorylation of SMO and other GPCRs is presently unclear.

The structural mechanism by which SMO phosphorylation enhances binding of PKA-C is also mysterious, as SMO is, to our knowledge, the first example of a PKA-C decoy substrate whose binding to PKA-C is regulated by phosphorylation^120–122^. One possibility is that GRK2 phosphorylation alters the conformation of the pCT to expose the otherwise “hidden” PKI motif. This model is unlikely because a peptide including only the SMO PKI motif (and no other SMO pCT sequences) binds PKA-C with only modest affinity (923 nM), similar to that of the nonphosphorylated SMO pCT (752 +/-34 nM)^39^. Instead, we favor a model based on avidity, in which GRK2 phosphorylation creates a new PKA-C binding site on SMO that synergizes with the modest-affinity PKI motif to enable efficient binding and inactivation of PKA-C^39^. Alternatively, SMO phosphorylation may alter the PKI motif’s conformation to increase its affinity for PKA-C, possibly via long-range allosteric interactions within the SMO pCT. Future structural studies of SMO-GRK complexes, as well as complexes between phosphorylated SMO and PKA-C, will help to address these questions.

### Kinetics of SMO phosphorylation suggest SMO and PKA-C regulatory mechanisms

We observed that SMO phosphorylation by GRK2, although occurring almost immediately after SMO activation, depends on continuous action of these kinases, as treatment with a GRK2 inhibitor triggers disappearance of phosphorylated SMO within minutes. The mechanisms underlying this rapid disappearance are currently unknown. Levels of SMO remain constant in these experiments, ruling out effects of phosphorylation on SMO stability. The rapid kinetics hint that one or more protein phosphatases^123,124^ may dephosphorylate SMO, either tonically or in response to SMO activation, thereby counterbalancing the effect of GRK2. Such a mechanism may serve to adjust SMO phosphorylation levels, potentially impacting the timing and intensity of SMO-GLI communication during Hh signal transduction. Future studies can evaluate these hypotheses by identifying candidate phosphatases that act on SMO and delineating their underlying modes of regulation. Our studies also reveal that PKA-C colocalizes with phosphorylated SMO, but with somewhat slower kinetics than those of SMO phosphorylation. Although these findings are consistent with SMO phosphorylation preceding and triggering PKA-C interaction, the mechanistic basis for the observed delay in PKA-C interaction remains unclear.

It is possible that this reflects a technical limitation — SMO phosphorylation may be easier to detect than PKA-C ciliary accumulation. Alternatively, PKA-C might not gain access to SMO in cilia via simple diffusion, but rather might be titrated via a regulated trafficking process that dictates the timing and extent of PKA-C ciliary localization. Such a mechanism would likely influence Hh along with other PKA-dependent ciliary pathways and may be revealed by more detailed studies of PKA-C ciliary trafficking mechanisms.

### GRK control of GPCR signaling in the cilium and beyond

Beyond SMO and GRK2 in the Hh pathway, our study has general implications for GRKs and for other ciliary GPCRs. Our work provides the first demonstration that a GRK can localize to the cilium, and that a ciliary GPCR undergoes GRK-mediated phosphorylation. It seems unlikely, however, that the pool of GRK2 at the base of the cilium is singularly dedicated to serving the needs of SMO during Hh signal transduction. Indeed, the cilium is home to a host of GPCRs critical to the nervous, cardiovascular, and musculoskeletal systems, and dysregulation of these receptors is associated with a range of devastating pathologies^9,10,125^. Many of these GPCRs are assumed to undergo agonist-dependent GRK phosphorylation in the cilium, but to our knowledge, this has never been demonstrated directly. We expect that the concepts and approaches developed here will enable studies of GRK ciliary localization as well as phosphorylation of ciliary GPCRs and the ensuing downstream signaling processes. Such studies will enable a better understanding of how GPCRs signal within cilia and how dysregulation of these processes instigates disease.

Whereas conventional plasma membrane GPCRs, require Gýy subunits for GRK2-mediated phosphorylation, we find that SMO phosphorylation in cilia is Gýy-independent, constituting the first example of a GPCR that undergoes GRK2 phosphorylation without participation from Gýy subunits. This finding raises the possibility that GRK2 recognizes GPCRs in the cilium via a mechanism distinct from that in the cell body. We speculate that in the highly confined environment of the primary cilium, other determinants of GPCR-GRK2 interactions may render contributions of Gýy unnecessary. This view is supported by prior findings that the GRK2 aN helix and membrane phosphoinositide-binding residues are necessary for SMO-GLI communication, whereas Gýy-binding residues are dispensable^44^. However, an important caveat is that the GRK2CT construct, while extensively characterized as a Gýy inhibitor in the plasma membrane^74–76^, has not yet been shown to block GRK2 phosphorylation or Gýy-dependent signaling for any ciliary GPCRs; thus, whether this construct can actually sequester Gýy subunits in the cilium remains to be demonstrated directly. In the future it will be worthwhile to use the methodologies we employed for SMO to study other GPCRs that undergo GRK2 phosphorylation in the cilium.

Lastly, GRK phosphorylation of GPCRs is canonically associated with β-arrestin binding^35,36,41,42^. In contrast, our studies reveal a new type of role for GRKs, in which receptor phosphorylation triggers direct interactions with PKA-C. This finding suggests that GRK phosphorylation may enable activated GPCRs to directly bind a variety of intracellular signaling factors, including β-arrestin, PKA-C, and perhaps other proteins. Such mechanisms may enable GPCR activation to encode a broad range of signaling outputs and thereby produce a wide array of biological outcomes. Given the emerging examples of direct GPCR coupling to factors other than heterotrimeric G proteins and β-arrestins^126–129^, understanding these signaling mechanisms may provide an exciting research direction in the coming years.

## ACKNOWLEDGMENTS

We thank M. Nachury and S. Nakielny for providing feedback on our manuscript, and K. Verhey for the gift of NIH3T3 Flp-in *Kif3a^-/-^* cells. This research project was supported in part by the University of Utah Cell Imaging Core Facility and the Emory University Integrated Cellular Imaging Core. This work was supported by an Ellison Foundation Research Scholar Grant from the American Cancer Society (R.S.), an NSF CAREER award (IOS-2143711) (X.G.) and the National Institutes of Health grant numbers R01HL071818 and R01CA254402 (J.J.G.T.), 1R01NS106527 (R.A.S.), 1R01GM145651 (R.S.), 1R15CA235749-01 (X.G.), 5R01GM143276-02 (X.G.), and 1R35GM133672 (B.R.M.). The funders had no role in study design, data collection and analysis, decision to publish, or preparation of the manuscript.

## SUPPLEMENTARY MOVIES

**Supplementary Movie 1.** Live-cilium TIRF movie of IMCD3 cells stably expressing GRK2-eGFP, with images acquired at 3 minute intervals. Cells are pre-imaged for 15 min, and SAG21k is added at the 0 min timepoint, followed by continued monitoring (see Main text). Labels and colors are as in Fig. 1. The SiR-Tubulin (red) and GRK2-eGFP (white) channels are shifted laterally with respect to one another, to facilitate viewing.

## SUPPLEMENTARY FIGURES

**Figure S1:**
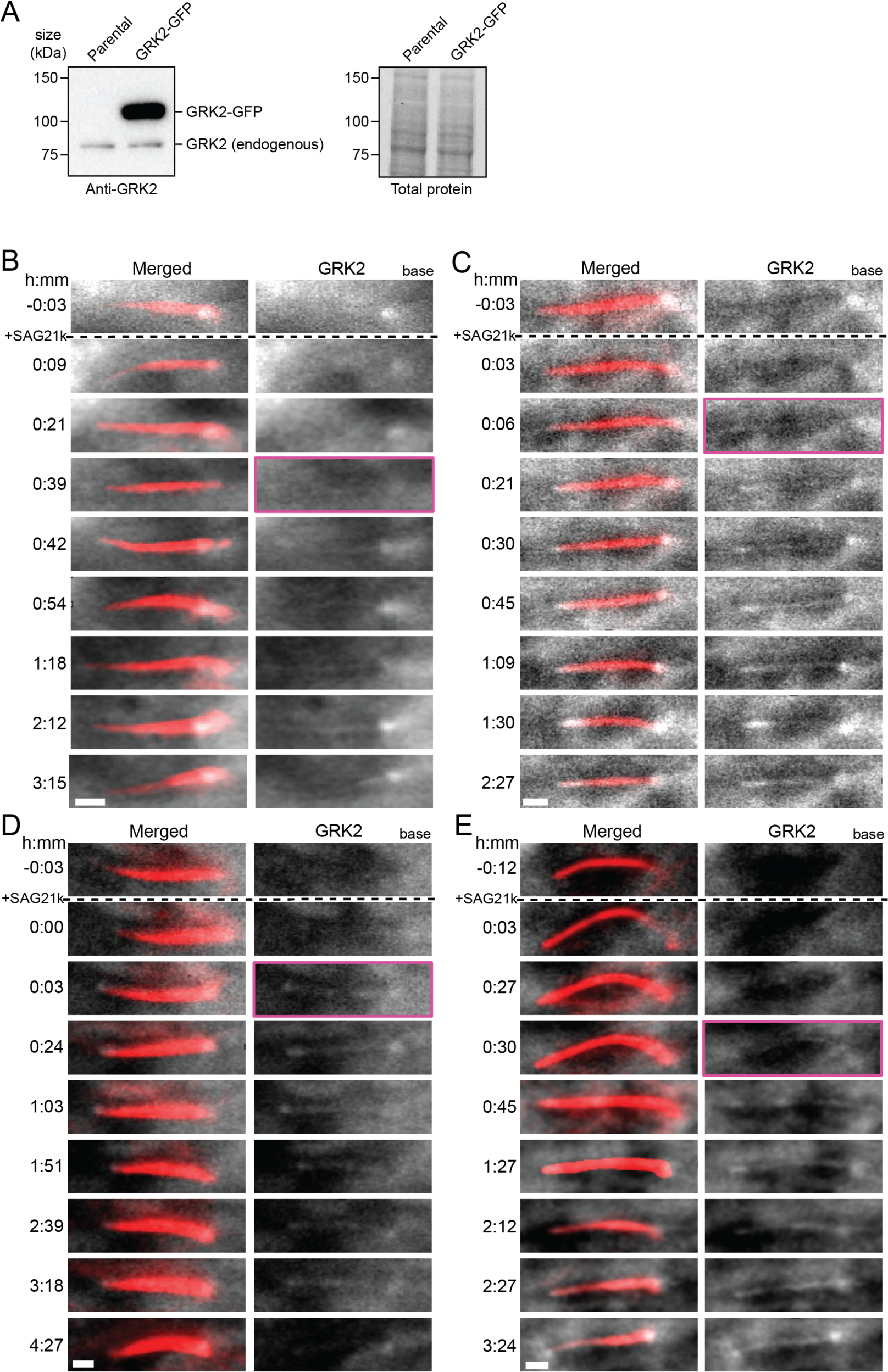
Characterization of IMCD3 cells stably expressing GRK2-eGFP, and additional representative TIRF images of GRK2-eGFP ciliary localization. **(A)** Expression of the GRK2-eGFP fusion was verified via immunoblot analysis in stably expressing IMCD3 Flp-in expressing the GRK2-eGFP fusion, revealing that the GRK2-eGFP was expressed at 23.99 +/-0.06-fold higher levels relative to endogenous GRK2. Note that the GRK2-eGFP fusion runs at a higher molecular mass, due to the eGFP tag. Left: anti-GRK2 immunoblot; Right: total protein (Stain Free imaging).(**B-E)** Four additional examples of cells showing SMO activation-induced localization of GRK2-eGFP to the ciliary shaft, monitored by TIRF imaging. Merged (SiR-Tubulin (red) to mark cilium and GRK2-eGFP (white) images) and GRK2-eGFP channels are shown, as in Fig. 1C.

**Figure S2:**
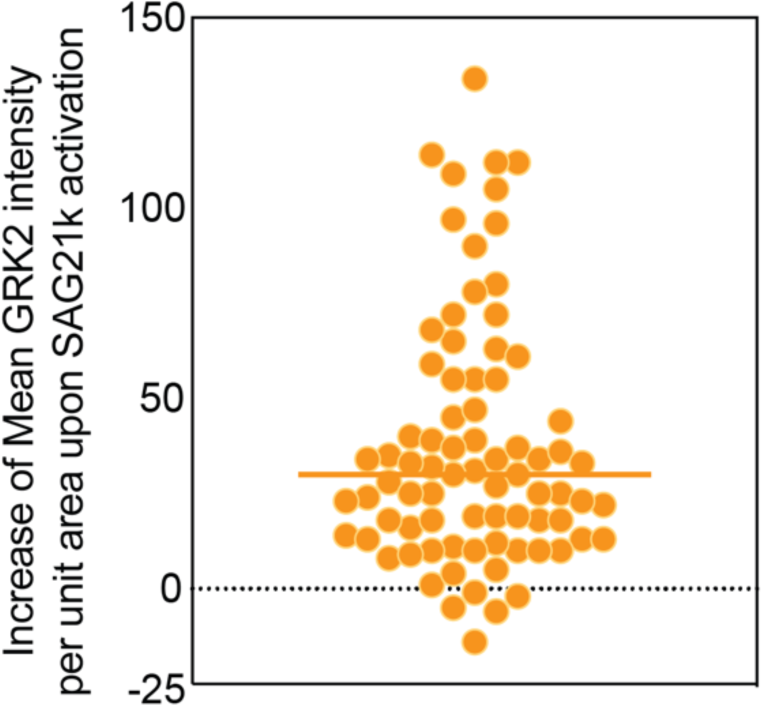
Increase of mean intensity of GRK2 per unit area in cilium shaft upon SAG21k activation. Scatterplot indicating the increase in mean GRK2 intensity per unit area within the cilium shaft following SAG21k activation. The dashed line in the plot represents the median value.

**Figure S3:.**
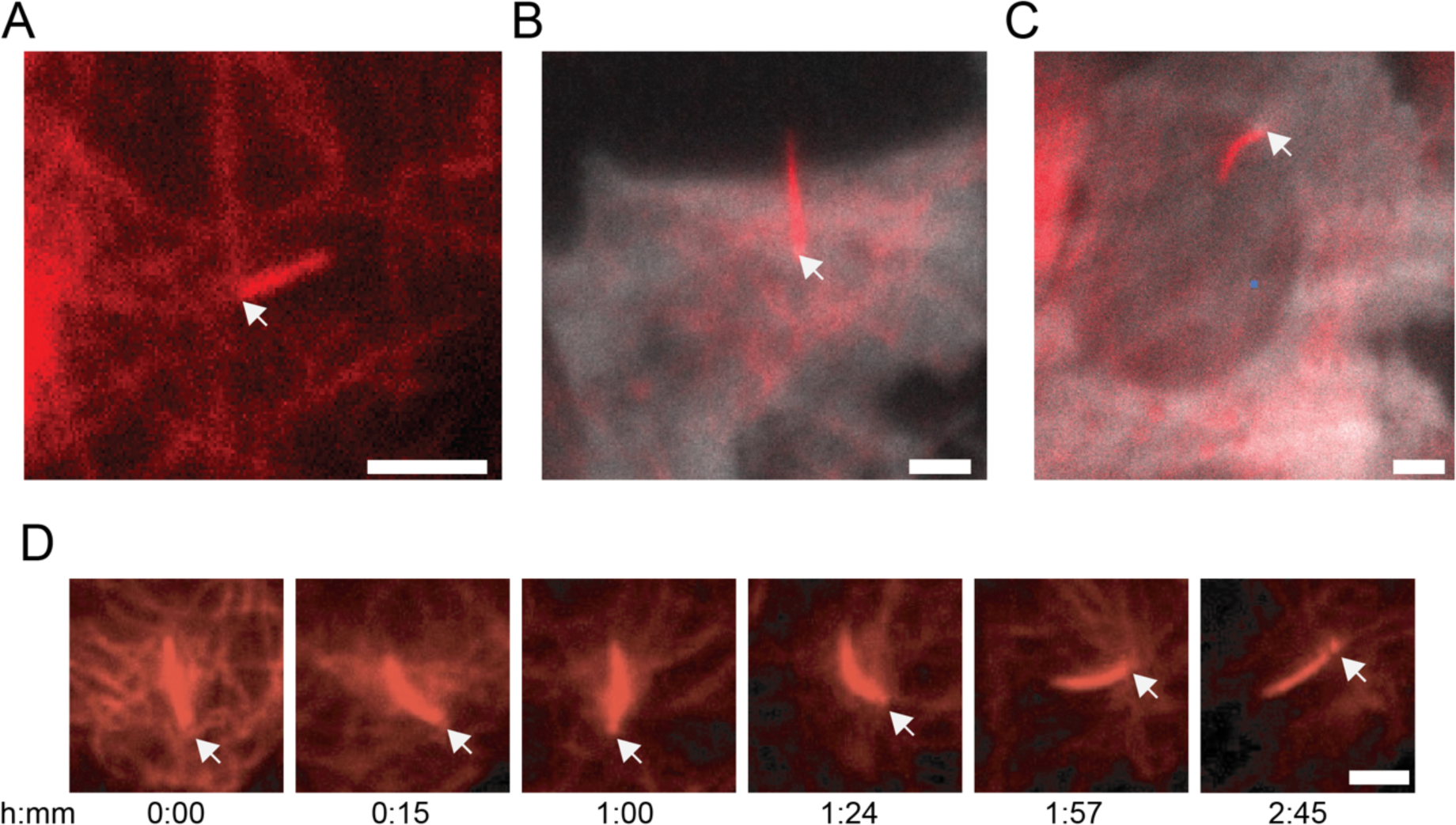
Determining the location of the cilium base. To distinguish the cilium base from the tip, several methodologies are employed. **(A)** The application of SiR-Tubulin dye, which stains both cytoplasmic microtubules and the ciliary axoneme, facilitates the visual identification of tubulin fibers converging from the cellular cytoplasm to the base of the cilia. In many instances, **(B)** cilia either extend from the cellular body into the extracellular space or **(C)** emanate from the nuclear periphery. **(D)** In cases where the determination of cilia base positioning is not apparent from an initial image, we followed the cilium over multiple timepoints; in many cases, such montages depict tubulin fibers converging towards the cilium base at later timepoints, enabling us to visualize the centriole (last image) and thereby distinguish the base from the tip. (Arrow points to the cilium base. Scale bar = 3μm).

**Figure S4:**
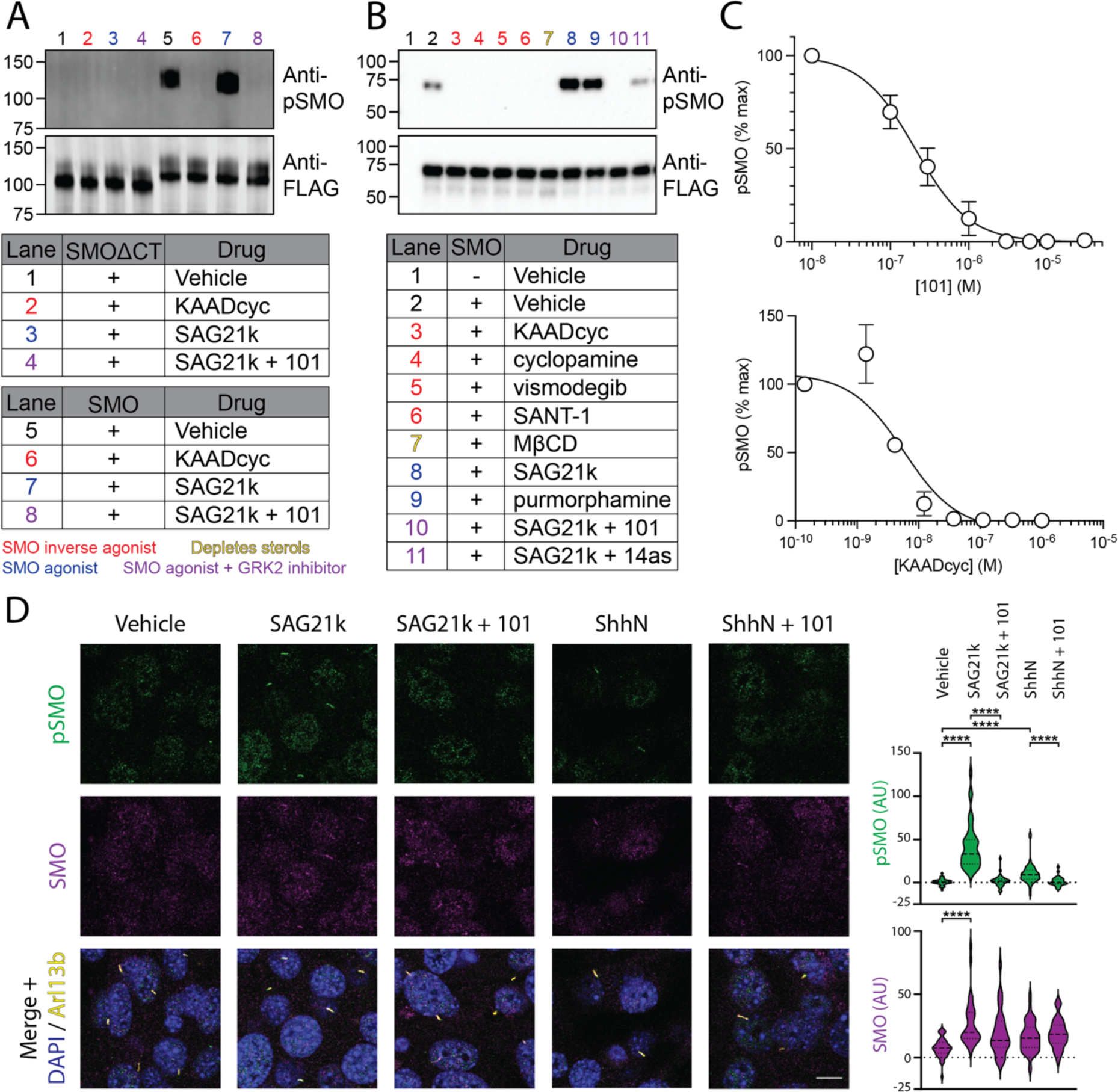
Validation of anti-pSMO antibody. **(A)** HEK293 cells expressing GRK2-GFP along with C-terminal Gα_o_ fusions of FLAG-tagged wild-type (WT) SMO or a SMO truncation mutant lacking all GRK2 phosphorylation sites (SMOΔCT) (see “Methods”) were treated for four hours with vehicle, the SMO agonist SAG21k (1 µM), the SMO inverse agonist KAADcyc (1 µM), or SAG21k plus the GRK2 inhibitor Compound101 (101, 30 µM). SMO was then purified using FLAG affinity chromatography, and FLAG eluates were analyzed via Western blot with anti-pSMO or anti-FLAG antibodies to probe phosphorylated SMO and total SMO, respectively. **(B**) HEK293 cells expressing GRK2-GFP and wild-type SMO (or a vector control) were treated with the indicated SMO inverse agonists (red), agonists (blue), agonist + GRK2 inhibitor (purple), or depleted of sterols using the cholesterol extracting agent methyl-β-cyclodextrin (MβCD, yellow), then analyzed as described in (A). Note that SMO overexpressed in HEK293 cells has substantial basal activity, as it exceeds regulation by endogenous PTCH1^105–107,130^ and is therefore constitutively bound to membrane sterols^27,107^. Consistent with this hypothesis, the anti-pSMO signal in vehicle-treated cells is absent in MβCD-treated cells. **(C)** Concentration-response analysis for blockade of SMO phosphorylation by KAADcyc or 101 revealed IC_50_ values of 5.8 nM and 207.7 nM, respectively, close to previously published values from Hh pathway transcriptional reporter assays^44,55^. **(D)** NIH3T3 cells were treated overnight with SAG21k (500 nM) or the N-terminal signaling domain of Sonic hedgehog (ShhN) (supplied as conditioned medium), in the presence or absence of 101 (30 µM), or with a vehicle control. Cells then were stained with anti-pSMO (green) to view phosphorylated SMO, anti-SMO (magenta) to view total SMO levels, and anti-Arl13b (yellow) to view primary cilia. Quantification of pSMO and SMO signals is shown at right. Significance in (D) was determined via a Mann-Whitney test. ****, p < 0.0001. n = 34-64 individual cilia per condition. Scale bar = 10 μm in all images.

**Figure S5:**
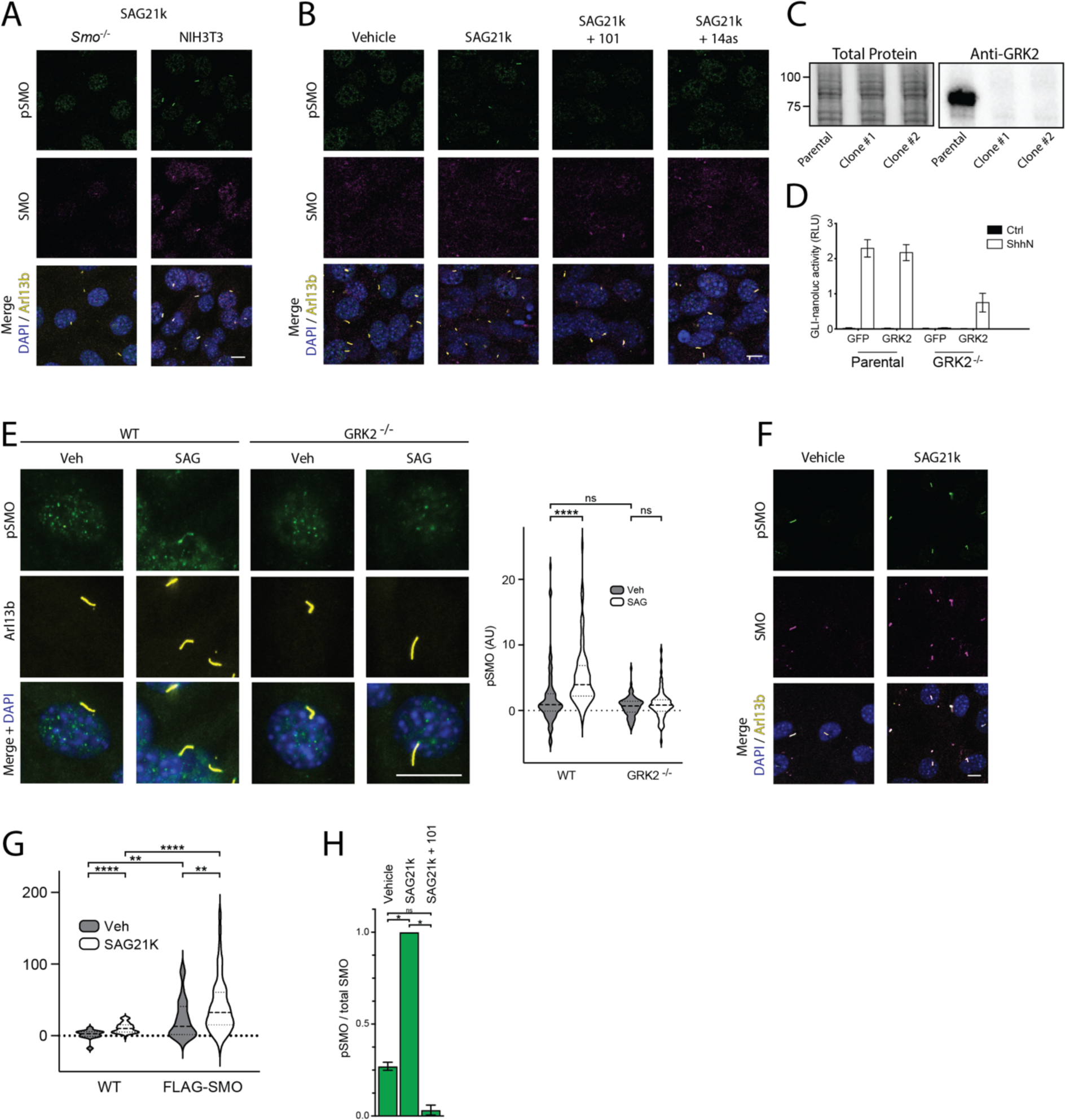
Additional validation of anti-pSMO antibody. **(A)** *Smo^-/-^* mouse embryonic fibroblasts (MEFs) or NIH3T3 cells were treated with SAG21k, then stained as described in Fig S4D. Note that the anti-pSMO antibody recognizes both a ciliary and a nuclear antigen. Only the latter is present in *Smo*^-/-^ MEFs, indicating that the nuclear signal is nonspecific and arises from a non-SMO antigen. **(B)** Effects of two independent GRK2 inhibitors, 101 and 14as, on SAG21k-induced SMO phosphorylation in NIH3T3 cells treated and stained as in (A). **(C)** Absence of GRK2 protein in NIH3T3 *Grk2^-/-^*cells was verified by immunoblotting. Lysates from two independent *Grk2^-/-^* clones are shown. **(D)** NIH3T3 parental or *Grk2^-/-^* cells were transfected with a GRK2 expression plasmid or a GFP control, and Hh pathway activity in response to ShhN (white) vs. control (black) conditioned medium was monitored via a GLI transcriptional reporter assay (see Methods). Note that *Grk2^-/-^* cells fail to respond to ShhN, and pathway responsiveness is restored when GRK2 (but not a GFP control) is reintroduced via transfection, consistent with previous findings^43,44^ **(E)** Wild-type or *Grk2^-/-^*NIH3T3 cells were treated for 4 hours with SAG vs. a vehicle control, then stained with anti-pSMO, Arl13b, and DAPI as in (A). **(F)** NIH3T3 cells stably expressing FLAG-SMO were treated with SAG21k or a vehicle control, then stained with anti-pSMO (green), anti-FLAG (to detect total SMO, magenta), and anti-Arl13b (yellow). Scale bar = 10 μm in all images. **(G)** Expression levels of the stably overexpressed SMO were quantifed relative to endogenous SMO via ciliary immunofluorescence staining in NIH3T3 Flp-in parental vs. FLAG-SMO-expressing cell lines with anti-SMO antibody, revealing that SMO is overexpressed 2.83-fold relative to the endogenous protein. **(H)** Quantification of the immunoblot data in Fig. 2D (mean +/-standard deviation from two independent experiments.) Significance was determined via one-way ANOVA. *, p < 0.05; ns, not significant.

**Figure S6:**
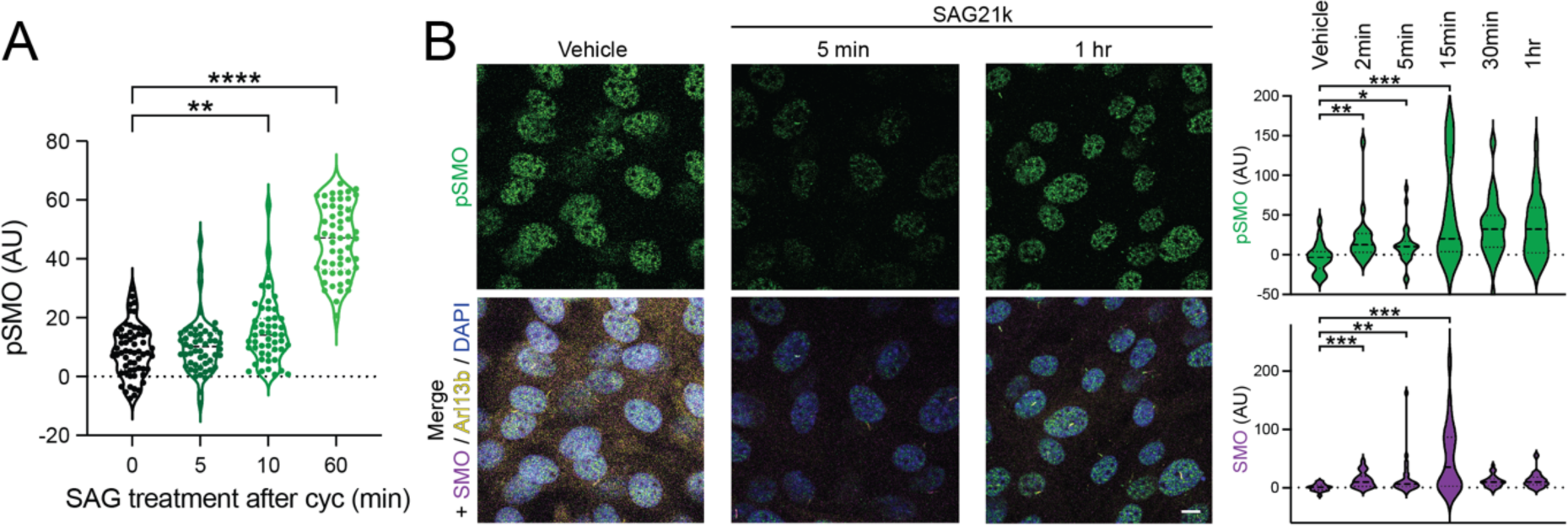
Additional time-course analysis of ciliary SMO phosphorylation in NIH3T3 cells. **(A)** Quantification of pSMO intensities in individual cilia from the NIH3T3 cells pretreated with cyclopamine and treated with SAG in Fig. 3A. **(B)** NIH3T3 dync2 cells were treated with SAG21k for the indicated times, then processed and stained as described in Fig. 2C. Quantification is shown at right. Note that the anti-pSMO antibody specifically recognizes phosphorylated SMO in cilia but also nonspecifically stains a nuclear antigen (see Fig. S5A). Significance in (A) was determined as in Fig. 3A, and in (B) was determined via a Mann-Whitney test. *, p < 0.05; **, p < 0.01; ***, p < 0.001; ****, p < 0.0001. n = 20-43 individual cilia per condition. Scale bar = 10 μm.

**Figure S7:**
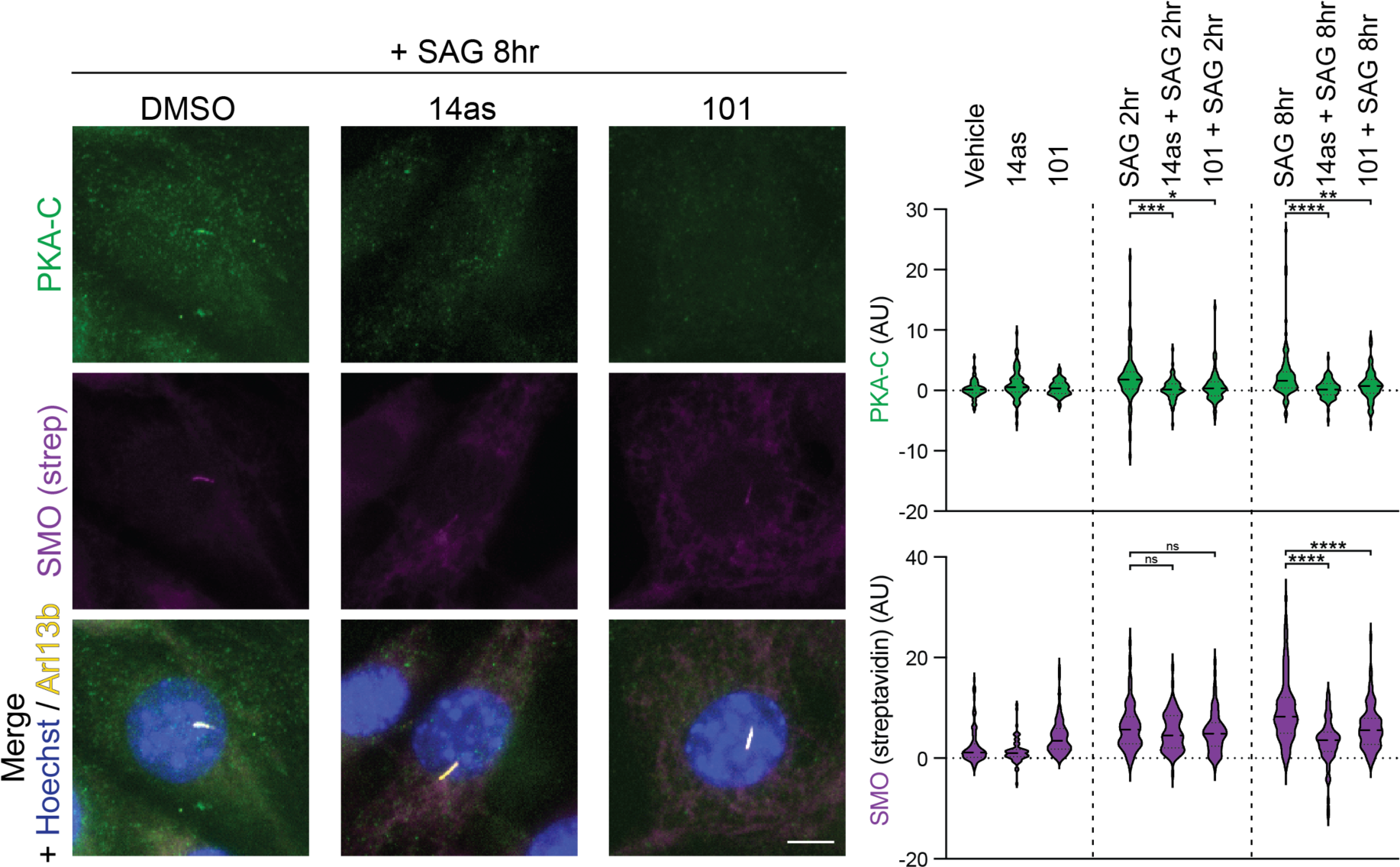
Additional timepoints for studies of GRK2 effects on SMO / PKA-C colocalization in cilia. The complete experiment from Fig. 5B, in which NIH3T3 cells stably expressing SMO-V5-TurboID were treated with vehicle, Cmpd101 or 14as for 16 hr. SAG was then added to the culture medium in the presence of the indicated inhibitors and incubated for 2 hr or 8 hr. Cells were labeled with biotin for 10 min before fixation. For simplicity, the 2 hr timepoint is presented in the main figure panel, and the complete experiment including all timepoints is presented here. Note that while the GRK2 inhibitors abolish ciliary SMO / PKA-C colocalization at 2 hr without affecting total SMO levels in cilia, these inhibitors did block the further accumulation of SMO in cilia between 2 hr and 8 hr, suggesting that prolonged GRK2 inhibition may affect SMO ciliary trafficking under some conditions. Such effects, however, cannot account for the effects of GRK2 inhibitors on SMO / PKA-C localization at the earlier timepoint, leading to the conclusion that these inhibitors block SMO / PKA-C interaction in cilia primarily by directly disrupting the SMO / PKA-C complex, rather than by affecting SMO ciliary accumulation. Significance was determined as in Fig. 5. *, p <0.05; **, p < 0.01; ***, p < 0.001; ****, p < 0.0001. n = 90-100 individual cilia per condition. Scale bar = 5 μm in all images.

**Figure S8:**
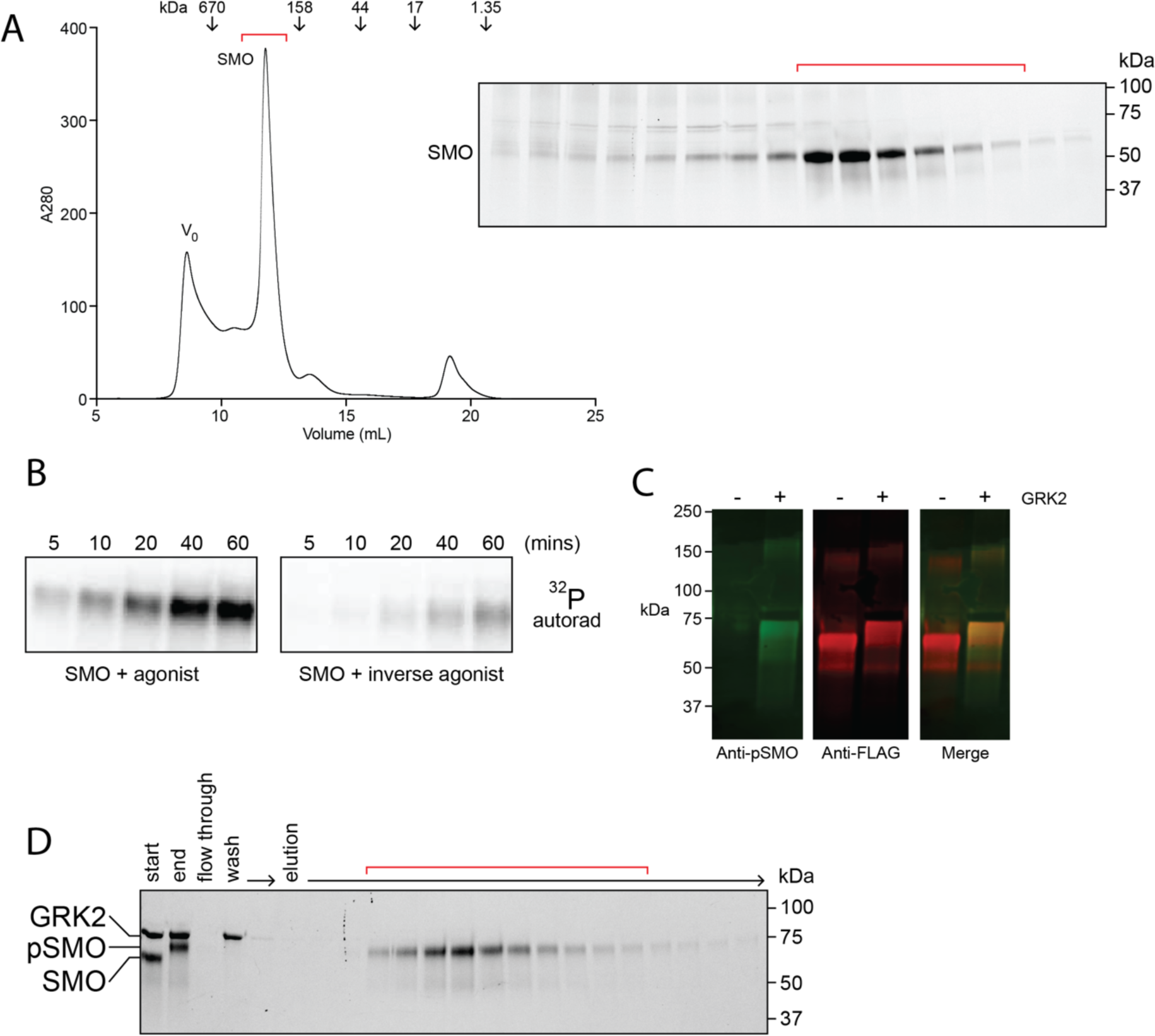
Purification of near-full-length SMO protein, and its phosphorylation by GRK2 *in vitro*. **(A)** Size exclusion chromatography of FLAG-SMO affinity-purified from HEK293 cells. Positions of the void (V_0_) and molecular weight standards are indicted. The peak corresponding to monodisperse SMO (red bracket) was collected and used for *in vitro* reconstitution studies. SDS-PAGE analysis of monodisperse SMO fractions is shown at right. **(B)** Phosphorylated agonist-or inverse-agonist loaded SMO was prepared as in Fig. 6B but in the presence of γ^32^P-ATP, then analyzed by autoradiography. **(C)** Immunoblot analysis of purified SMO before and after the GRK2 phosphorylation reaction, demonstrating phosphorylation at the GRK2 cluster recognized by our anti-pSMO antibody. **(D)** Following GRK2 phosphorylation, pSMO was separated from GRK2 via FLAG affinity chromatography. Flow-through, wash, and elution fractions from the purification procedure are analyzed by SDS-PAGE. “Start” and “end” correspond to GRK2 phosphorylation reaction prior to addition of ATP (“start”) or after completion of the reaction (“end”). Red brackets indicate elution fractions that were collected, pooled, and used for subsequent experiments. Positions of SMO, pSMO, and GRK2 are indicated at left.

**Figure S9:**
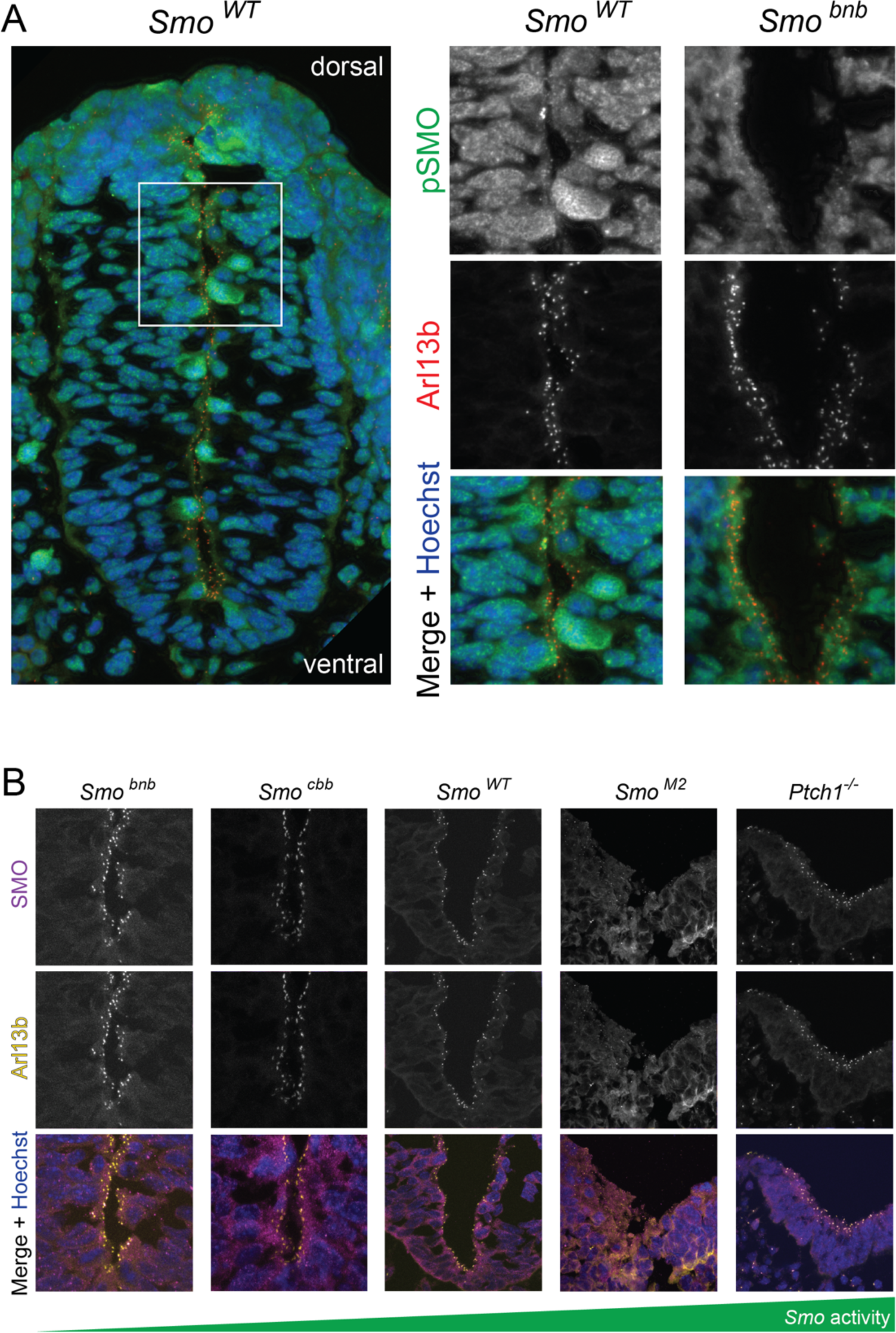
Ciliary SMO phosphorylation in the mouse dorsal neural tube. **(A)** Neural tubes from the indicated wild-type or *Smo^bnb^* mutant E9.5 mice (low-magnification view of wild-type mouse at left, higher-magnification view at right, boxed region indicates the zoomed-in portion) were stained for pSMO and Arl13b, as in Fig. 7A. Images are oriented with dorsal pointing up and ventral pointing down. (**B)** Neural tubes from the indicated genotypes were stained for total SMO (magenta), Arl13b (yellow), and nuclei (Hoechst, blue).

**Figure S10:**
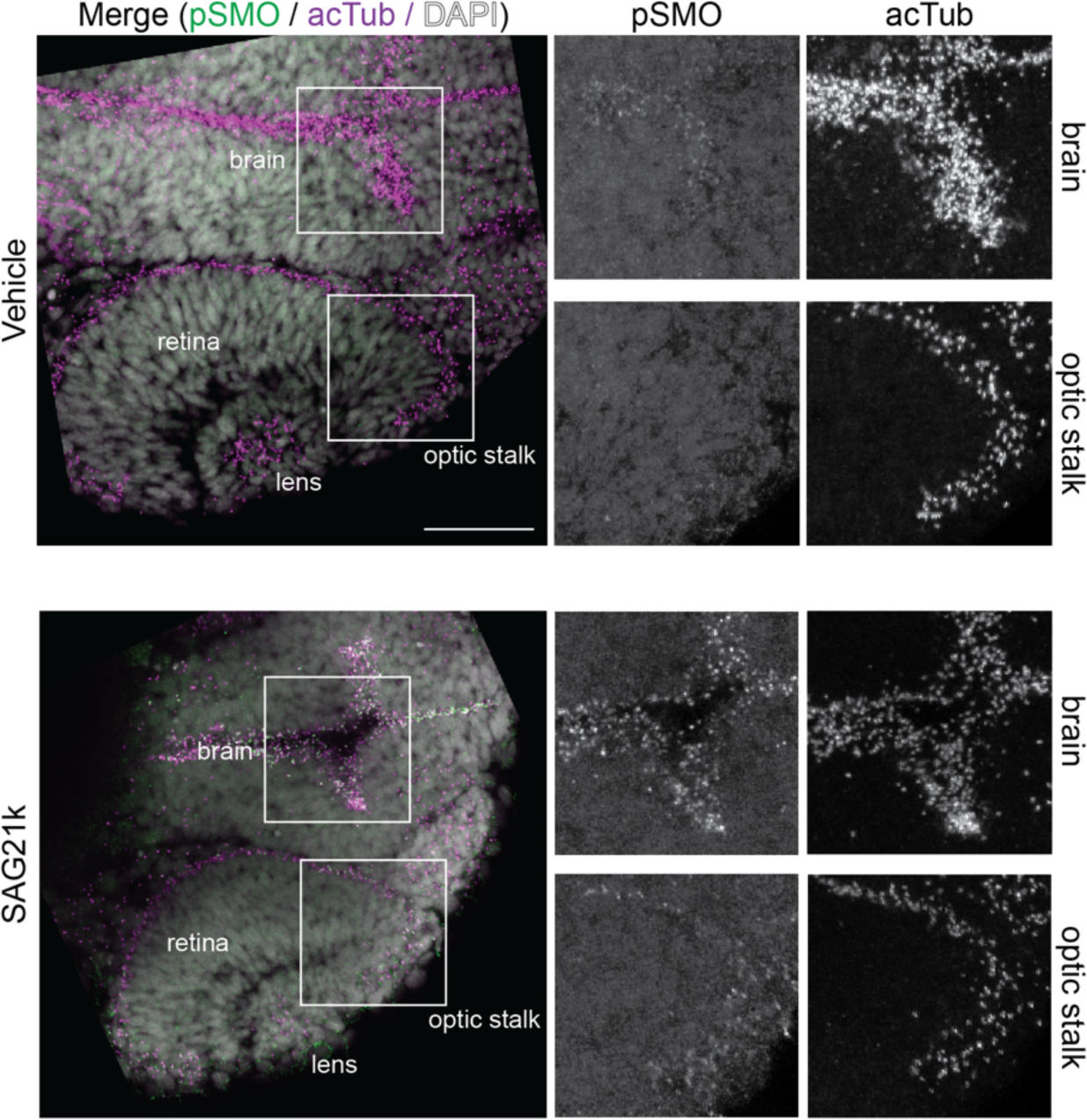
Ciliary SMO phosphorylation during zebrafish eye and forebrain development. Vehicle (DMSO) or SAG21k-treated (10 μM) zebrafish embryos were stained for pSmo (green), acetylated tubulin (acTub, magenta), and nuclei (DAPI, white). Locations of brain and lens are indicated with boxes. Merged image is shown at left, and zoomed-in views of individual channels within the boxed regions are shown at right. pSmo sparsely labels acetylated tubulin-positive punctae in vehicle-treated embryos, while SAG21k-treated embryos show extensive cilia labeling. All embryos are 24 hours post-fertilization, dorsal view. Scale bar, 50 µm.

**Figure S11:**
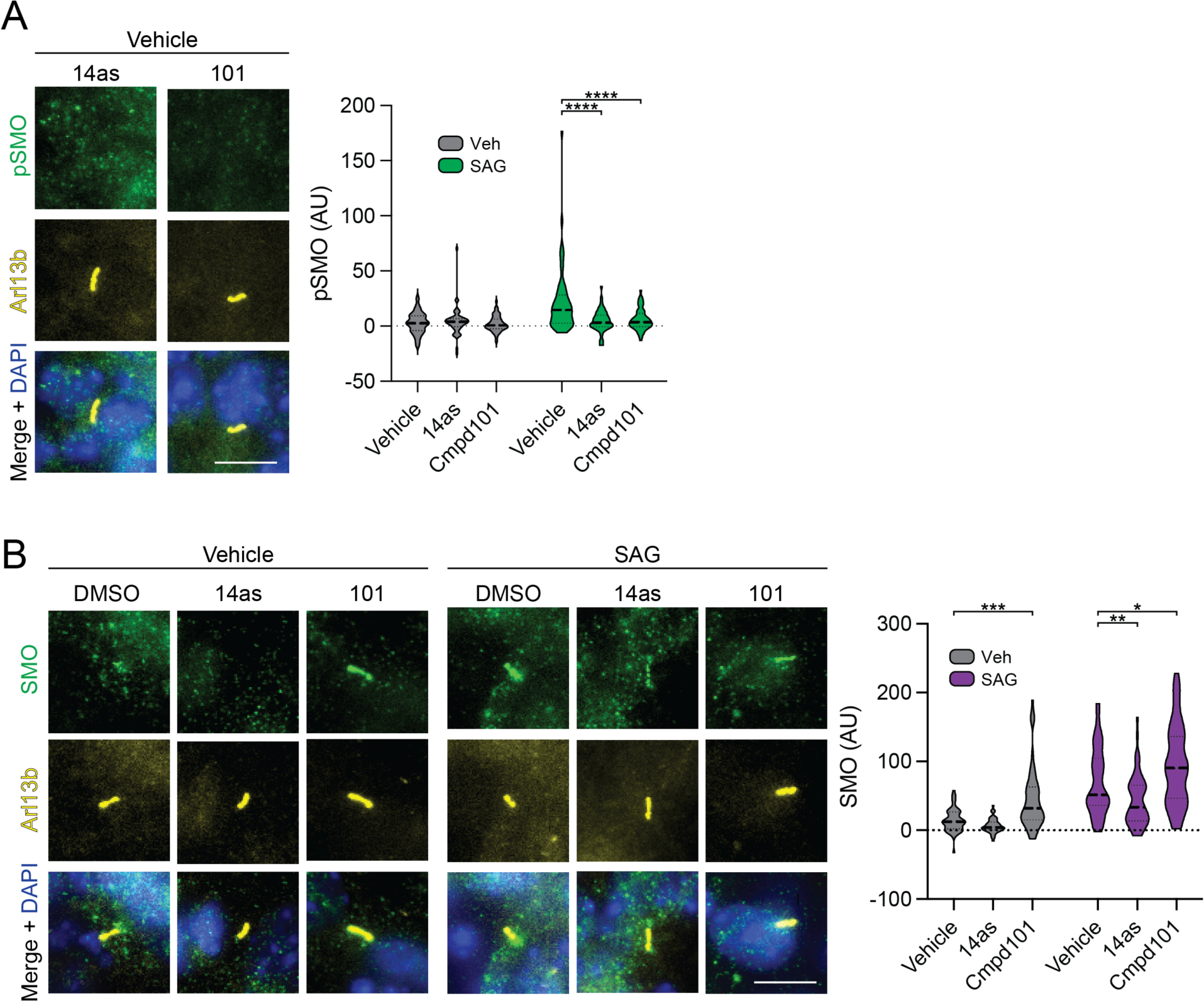
Additional data for experiments in cerebellar granule neural precursors (CGNP). **(A)** Additional raw images from the experiment in Fig. 7C, in which cells were treated with 14as or Cmpd101 in the absence of SAG, then stained for pSMO as described in the main figure. Quantification of the complete experiment is shown at right **(B)** CGNPs treated as in Fig. 7C were stained for total SMO (green), Arl13b (yellow), and DAPI (blue). Quantification is shown at right. *, p <0.05; **, p < 0.01; ***, p < 0.001; ****, p < 0.0001. Scale bar = 10 μm.

## MATERIALS AND METHODS

### Supplemental info on experimental model systems

#### Expression levels of GRK2-eGFP in stably transfected IMCD3 cell lines

Based on immunoblotting analysis of whole cell lysates from IMCD3 Flp-in parental and GRK2-eGFP-expressing cell lines, the GRK2-eGFP transgene is expressed at levels 23.99 +/-0.06-fold greater than endogenous GRK2 (Fig. S1A). Nevertheless, our findings regarding GRK2 ciliary localization are unlikely to be overexpression artifacts, because (1) this localization is not static, but rather changes in response to SMO activation (i.e., GRK2 relocalizes from the base to the shaft rapidly following addition of SMO agonist); (2) an earlier study^131^ detected endogenous GRK2 at the centrosome via immunofluorescence microscopy in a variety of cell lines, consistent with our finding that GRK2 localizes to the base of the cilium.

### Small Molecules and Biochemicals

SiR-Tubulin was obtained from SPIROCHROME (SC002). Compound101 was obtained from Hello Bio (HB2840). SAG21k was a gift from P. Beachy. SAG was obtained from Selleckchem (S7779). Cyclopamine was obtained from LC Laboratories (C-8700). KAADcyc was obtained from Toronto Research Chemicals (K171000). Vismodegib was obtained from LC Laboratories (V-4050). SANT-1 was obtained from ApexBio (A8659). Methyl-beta-cyclodextrin was obtained from Sigma Aldrich (332615). Purmorphamine was obtained from Cayman Chemicals (10009634). DAPI was obtained from Thermo-Fisher Scientific (D21490). Hoechst 33342 was obtained from Thermo-Fisher Scientific (H3570). SlowFade Gold was obtained from Thermo-Fisher Scientific (S36937). Prolong Gold was obtained from Thermo-Fisher Scientific (P36934). Fluoromount G was obtained from SouthernBiotech (0100-01). 14as (also referred to as GRK2-IN-1) was obtained from MedChem Express (HY-109562A). D4476 was obtained from MedChem Express (HY-10324). ATPγ32P, 3000 Ci/mol was obtained from PerkinElmer (BLU502H250UC).

### Antibodies

Mouse anti-SMO (E-5) was obtained from Santa-Cruz Biotechnology (sc-166685). Rat anti-Arl13b was obtained from BiCell Scientific (90413). Mouse anti-Arl13b (clone N295B/66) was obtained from Antibodies Inc (75-287). Rabbit anti-Arl13b was obtained from Proteintech (17711-1-AP). Rabbit anti-SMO antibodies were previously described.^20,31^ Rabbit anti-pSMO was obtained from 7TM antibodies (7TM0239A). Mouse anti-PKA-C was obtained from BD Biosciences (610980). Mouse anti-GRK2 (C-9) was from Santa Cruz (sc-13143). Alexa 555 anti-rat was obtained from Thermo-Fisher Scientific (A21434). Alexa 488 anti-rat was obtained from Thermo-Fisher Scientific (A11006). Alexa 488 anti-rabbit was obtained from Thermo-Fisher Scientific (A32731). Alexa 555 anti-rabbit was obtained from Thermo-Fisher Scientific (A32732). Alexa 647 anti-mouse was obtained from Thermo-Fisher Scientific (A32728). Alexa 568 anti-mouse was obtained from Thermo-Fisher Scientific (A10037). Anti-Mouse IgG (H+L), HRP Conjugate was obtained from Promega (W4021). Anti-Rabbit IgG (H+L), HRP Conjugate was obtained from Promega (W4011). Goat Anti-Rabbit IgG Antibody, IRDye 800CW Conjugated was obtained from LiCor Biosciences (926-32211). Goat Anti-Mouse IgG Antibody, IRDye 680RD Conjugated was obtained from LiCor Biosciences (926-68070). Mouse anti-FLAG M2 affinity gel was obtained from Sigma Aldrich (A2220-1ML). Mouse anti-FLAG M1 was made in-house.

### Biochemistry and Protein Analysis

4-20% Crit TGX Stain-Free Gel 26W was obtained from Bio-Rad Labs (5678095). 4-20% Crit TGX Stain-Free Gel 18W was obtained from Bio-Rad Labs (5678094). Non-Fat Dry Milk was obtained from Lab Scientific (M-0841). Laemmli Sample Buffer was obtained from Bio-Rad Labs (1610737EDU). Immobilon-P PVDF Membrane was obtained from EMD Millipore (IPVH00010). Pierce™ Protease Inhibitor, EDTA Free was obtained from Thermo-Fisher Scientific (A32965). DDM (n-Dodecyl-β-D-maltoside) Anagrade, ≥ 99% β+α was obtained from Anatrace (D310 5 GM). CHS (Cholesteryl Hemisuccinate Tris Salt) was obtained from Anatrace (CH210 5 GM). GDN (glyco-diosgenin) was obtained from Anatrace (GDN101 5 GM). CNBr-Activated Sepharose 4 Fast Flow was obtained from GE Healthcare (17-0981-01). NanoGlo lytic detecton system was obtained from Promega (N3030). Triton X-100 was obtained from Sigma-Aldrich (X100-100ML). Bovine Serum Albumin was obtained from Sigma-Aldrich (A4503-50G). Tween 20 was obtained from Sigma-Aldrich (P1379-500ML). PBS was obtained from Thermo-Fisher Scientific (10010049). PFA was obtained from Fisher Scientific (50-980-495).

### Cell Culture Reagents

Papain was obtained from Worthington (LS003126). Dnase I was obtained from Roche (11284932001). Neurobasal medium was obtained from Gibco (21103049). B-27 Supplement was obtained from Gibco (17504044). GlutaMAX Supplement was obtained from Gibco (35050061). Pen Strep was obtained from Gibco (15140122). Laminin was obtained from Gibco (23017015). DMEM / F12 medium was obtained from Gibco (11330-057). Trypsin EDTA .05% was obtained from Life Technologies (25300-120). DMEM, High Glucose, GlutaMAX Supplement, Pyruvate was obtained from Thermo-Fisher Scientific (10569044). DMEM, high glucose was obtained from Thermo-Fisher Scientific (11965118). Fetal Bovine Serum was obtained from Omega Scientific (FB-02). Calf Serum (HyClone) was obtained from Fisher Scientific (SH3007203). Sf-900™ III SFM was obtained from Thermo-Fisher Scientific (12658019). Gentimicin (10mg/mL) was obtained from Thermo-Fisher Scientific (15710064). Gibco™ FreeStyle™ 293 Expression Medium was obtained from Thermo-Fisher Scientific (12-338-018). Penicillin-Streptomycin-Glutamine (100X) was obtained from Life Technologies (10378016). BestBac 2.0 Δ v-cath/chiA Linearized Baculovirus DNA was obtained from Expression Systems (91-002).

### Transfection Reagents

Fugene 6 was obtained from Promega (E2691). Lipofectamine 2000 was obtained from Thermo-Fisher Scientific (11668019). Cellfectin™ II was obtained from Thermo-Fisher Scientific (10362100).

### DNA constructs

HEK293 cell studies employed either: 1) FLAG-SMO, an N-terminally FLAG-tagged construct containing the mouse SMO (mSMO) extracellular cysteine-rich domain (CRD), 7TM domain, and pCT domains (truncated at mSMO residue 674, to remove the unstructured distal C-tail (dCT) domain); or 2) FLAG-SMOΔCT, which lacks the entire intracellular domain (truncated at mSMO residue 666). Initial experiments characterizing our anti-pSMO antibody on purified SMO proteins (Fig. S4A) employed C-terminal Gɑ_o_ fusions of these constructs (FLAG-SMO655-Gɑ_o_ or FLAG-SMO566-Gɑ_o_), which increases biochemical stability of SMO in some settings^27,107^; however, we found in later experiments that the fused Gɑ_o_ was not necessary for SMO stability, and thus the majority of the experiments in this paper used a SMO construct lacking the C-terminal Gɑ_o_ fusion. SMO-V5-turboID in pEF5-FRT/hygro and mouse PKA-C in pRSET-B were previously described^39^. N-terminally His-tagged GRK2 / pFastBac was previously described^132^. FLAG-SMO-nanoluc-IRES-mNG3k/pEF5-FRT-hygro was constructed by fusing in tandem: 1) a full-length FLAG-tagged SMO construct containing the CRD, 7TM domain, pCT, and dCT, and with nanoluciferase (nanoluc) fused at the C-terminus; the native SMO signal sequence was replaced by an N-terminal FLAG tag and an HA signal sequence^38^; 2) an IRES element; and 3) a mNeonGreen3k fragment (mNG3k)^133^. The mNG3k was included to enable potential mNeonGreen labeling of endogenous NIH3T3 proteins that had been tagged with an mNG11 helix, via mNeonGreen complementation^133^; however, we ultimately did not pursue this strategy in our experiments, and so this feature of our expression vector was not used. Bovine GRK2-eGFP/pEF5-FRT was constructed by moving the GRK2-eGFP fusion from pVLAD6 to pEF5-FRT. GRK2CT was targeted to the cilium as previously described^52^, with minor modifications. Briefly, the ciliary localization signal from fibrocystin (MLSCVLCC…RMKV) was fused to mNeonGreen and the human GRK2 C-terminus (GIKLLDS…SANGL), and cloned into pEF5-FRT-hygro for stable transfection in NIH3T3 Flp-in cells; the negative control construct lacked the GRK2CT sequence. All DNA constructs were prepared in-house via Gibson assembly, or commercially (Epoch Life Sciences; Missouri City, TX), and verified by Sanger and/or next-generation sequencing before use.

### Cell culture and transfections

HEK293S-GnTI^-^ cells were grown in Freestyle 293 Expression Medium with 1% Fetal Bovine Serum, as previously described^38,39^. NIH3T3 Flp-In cells and NIH3T3 *dync2* cells^32^ were grown in DMEM with GLUTAMAX and 10% Calf Serum (HyClone) with Penicillin/Streptomycin (Thermo Fisher); to induce ciliation, the same medium but with 0.5% Calf Serum was used. IMCD3 cells stably expressing GRK2-eGFP were produced via stable transfection and selection in blasticidin as previously described^38,39^, and cultured in DMEM/F12 medium, supplemented with 10% FBS containing 1 mM sodium pyruvate and 2 mM L-glutamine, and were maintained in 5% CO_2_ at 37°C in a T-75 flask. Levels of GRK2-eGFP expression were quantified by anti-GRK2 immunoblotting. The NIH3T3 SMO-V5-turboID cell line was cultured as previously described^39^. The NIH3T3 Flp-In FLAG-SMO-nanoluc-IRES-mNG3k cell line was constructed by transfecting 2 µg of FLAG-SMO DNA + pOG44 into 70% confluent NIH3T3 Flp-in cells using Lipofectamine 2000, according to the manufacturer’s instructions. Cells were left to grow in antibiotic-free medium for 48 hours and then selected with Blasticidin (10 µg/mL) for 2 weeks. To identify clones expressing low levels of SMO, we isolated individual clones and screened for expression using the C-terminal nanoluc fusion (Nanoluc Lytic Detection Kit, Promega), followed by additional characterization of SMO agonist-induced ciliary accumulation via microscopy. Levels of FLAG-SMO expression were quantified by anti-SMO immunofluorescence microscopy. Conditioned medium containing the N-terminal signaling domain of Sonic hedgehog (ShhN), or control, non ShhN-containing conditioned medium, were prepared as previously described^56,134^. NIH3T3 Flp-in *Kif3a*^-/-^ cells were a gift from K. Verhey and were maintained and stably transfected as described above for NIH3T3 Flp-in cells.

### GLI reporter assays

GLI reporter assays in NIH3T3 Flp-in parental or *Grk2^-/-^*cells were performed as previously described, except that we reformatted the dual luciferase assay to utilize nanolucifrase and firefly luciferases. Briefly, the firefly luciferase in the 8xGli-firefly/pGl3b construct was replaced with nanoluciferase to make 8xGli-nanoluc/pGl3b. Cells were transfected in 24 well plates with 250 ng total DNA per well, 125 ng of which was the reporter plasmid mixture (1:1 ratio of 8xGli-nanoluc/pGl3b to pGL4.53[luc2/PGK] (Promega)), and 125 ng of which was GFP/pCDNA3. Cells were lysed in 100 μl of passive lysis buffer, and 25 μl of lysate was analyzed using the protocol described in the Nano-Glo® Dual-Luciferase® Reporter Assay System (Promega).

### Immunoprecipitation and immunoblotting

HEK293 GnTI^-^ cells were infected in 3 ml volumes with GRK2-GFP along with FLAG-SMO-674 or FLAG-SMO-566 BacMam viruses, and FLAG-tagged SMO was purified from lysates via M1 FLAG affinity chromatography as previously described, and blotted with anti-M1 FLAG (1:1000) or anti-pSMO (1:1000). NIH3T3 cells expressing FLAG-SMO-nanoluc (see above) were grown to confluency in 10 cm dishes (one 10 cm dish per experimental condition or timepoint). Once confluent, cells were moved to low-serum medium overnight, then treated with drugs for 4 hours. In the dephosphorylation experiment, cells were treated with SAG21k for 4 hours and then replaced with SAG21k/cmpd101 for the indicated times. Following treatment, cells were lysed in RIPA buffer with phosphatase and protease inhibitors. Lysates were centrifuged for 30 min at 4°C at 20,000 x g, and FLAG-tagged proteins purified using M2 FLAG resin. Following resin isolation and washing, bound proteins were eluted with 2x Laemmli Sample Buffer and samples were then separated via SDS-PAGE using 4-20% Criterion TGX gels (BioRad) and transferred to a PVDF membrane. The membrane was blocked with 5% Fat-Free Milk (or 2% BSA) and probed with anti-FLAG and anti-pSMO followed by HRP conjugate for chemiluminescence or IRdye conjugate for LiCOR imaging.

### Immunofluorescence microscopy of cultured fibroblasts

NIH3T3 Flp-In cells were seeded on glass or poly-D-lysine-coated coverslips and grown in DMEM GLUTAMAX + 10% Calf Serum. Once confluent, the media was then changed to low-serum medium (overnight) to allow ciliogenesis, along with respective drug treatments or vehicle controls. To activate Hh signaling, cells were stimulated with the N-terminal signaling domain of Sonic hedgehog (ShhN), the specific SMO agonist SAG, or the higher-affinity SAG derivative SAG21k. For time course studies, cells were serum starved in low-serum medium overnight to induce ciliogenesis, then treated for the indicated times with the indicated SMO agonist. The cyclopamine pretreatment was done by serum starving the cells in DMEM GLUTAMAX with 0.5% Calf Serum overnight with (5 µM) cyclopamine, followed by treatment with 100 nM SAG for the indicated times. For experiments with D4476, cells were either 1) treated overnight with the inhibitor (10 µM) dissolved directly in low-serum medium; or 2) to enhance cell permeability of D4476, a previously described cationic lipid complexation procedure was followed^61^ in which 3 µL of 100 mM D4476 was mixed with 9 µL of Fugene 6 and 18 uL of serum free DMEM, then allowed to incubate at 37°C for 15 minutes, then applied to 3 mL of DMEM + GLUTAMAX with 0.5% Calf Serum over the cells (100 µM), and incubated for 2 hr prior to fixation and staining. Similar results were obtained using both protocols 1 and 2; the data from protocol 1 is presented in the manuscript.

<u>For experiments in</u> Fig 2 and 4: Following incubation in low-serum medium, cells were washed with PBS, fixed in 4% PFA in PBS for 10 minutes, permeabilized in 0.1% Triton X-100 in PBS for 10 min, and blocked in 2% Bovine Serum Albumin and 1x Tris-Buffered Saline and 0.1% Tween-20 (TBST, 30 min at room temperature, or overnight at 4°C). Coverslips were then incubated at room temperature with primary antibodies diluted 1:1000 in TBST for 1 hour, followed by 3x 5 min washes in TBST. Coverslips then were incubated with the appropriate secondary antibodies diluted 1:1000 followed by 3x 5 min washes with TBST, then one wash with water. Coverslips were mounted onto a slide with SlowFade mounting medium. Images were acquired on a Leica SP8 laser scanning confocal using a 40x water immersion lens. For most experiments, we used Alexa 647-conjugated antibodies to visualize SMO, 555-conjugated antibodies to visualize Arl13b, and 488-conjugated antibodies to visualize pSMO. However, NIH3T3 *dync2* cells^32^ are engineered to stably coexpress GFP as a marker for the dync2 shRNA construct, creating significant background in the 488 channel. To mitigate this issue, GFP was denatured using a methanol fixation step (5 min, -20°C) following the PFA fixation step, and Arl13b was analyzed in the 488 channel while pSMO was analyzed in the 555 channel.

<u>For experiments in</u> Fig 3<u>, 5, and 7C:</u> cells were blocked with blocking buffer (0.2% Triton X-100, 2% Donkey serum in PBS) for 1 h at room temperature. After blocking, cells were incubated with primary antibody at 4°C overnight. Primary antibodies used were: rabbit anti-pSMO (1: 1000), rat anti-ARL13B (1:500), rabbit anti-ARL13B (1:1000), and mouse anti-PKA-C (1:200). Subsequently, cells were incubated with secondary antibodies for 1 hour and Hoechst 33342 for 10 min at room temperature. Cells were mounted in Fluoromount-G. Imaging was performed with a Zeiss LSM 880 confocal Laser Scanning Microscope with 100x oil immersion lens or a LEICA DMi8 system with ×63 oil-immersion lens.

For all immunofluorescence images, quantification of ciliary pSMO, SMO, or PKA-C intensity staining was performed in Fiji. Briefly, the contour of the cilium was outlined in the Arl13b channel, and the intensity within the area of interest was measured in the SMO, PKA-C or pSMO channels. After that, the contour was dragged to the area immediately adjacent to the cilium to measure the background values, which were then subtracted from the cilium intensity. The intensities are reported in Arbitrary units (AU). To quantify level of SMO overexpression in stable NIH3T3 Flp-in cells (Fig. 2D), parental vs stable cells were treated overnight with SAG21k, then stained for SMO (rabbit) and Arl13b (mouse) as described above. Levels of SMO in 56-66 individual cilia from parental cells (SMO_endo_ = 10.87) or overexpressing cells (SMO_FLAG =_ 41.6) were quantified as described above. The overexpression ratio was obtained by calculating: (SMO_FLAG_ – SMO_endo_) / SMO_endo_.

### Phospho-specific SMO antibody

The phosphorylation state-specific rabbit SMO antibody targeting pS594/pT597/pS599-SMO (7TM0239A) was provided by 7TM Antibodies (www.7tmantibodies.com).

### TIRF microscopy

For live-cell Total Internal Reflection Fluorescence (TIRF) microscopy, 1.5×10^5^ cells were seeded in a 35 mm glass bottom coverslip pre-coated with 0.1% gelatin in 1x PBS buffer. The cells were incubated in 5% CO_2_ at 37°C for 24 hours before transferring to low-serum DMEM (0.2% FBS) for at least another 24 hours to induce ciliogenesis. On the day of microscopy, cells were stained with 1 µM SiR-Tubulin and incubated for 1 hour, followed by three washes with low-serum DMEM to remove excess stain. We employed a TIRF microscope (Nikon) for live cell imaging (3 minutes per frame). Cells were excited with a 488 nm and a 647 nm laser for imaging eGFP and SiR-Tubulin, respectively. The images were captured on an Andor ZYLA CMOS camera controlled by the NIS-Elements software. A 100x objective lens (Nikon) with a NA = 1.45 was used to select cells where the cilium was attached to the coverslip surface. We identified the cilium using SiR-Tubulin staining, using the cytoplasmic eGFP signal to identify the orientation of the cilium and distinguish the base from the tip. Only cilia that were stably attached to coverslip for 15 minutes and showed detectable GRK2-eGFP localization at the base were used for subsequent SMO agonist (SAG21k) treatment and time-course analysis. The microscope chamber during imaging was maintained at 37°C and 5% CO_2_ in a designated dark room. Fluorescence intensity profile along the cilium were determined using line scan analysis in FIJI. Only the cilia where the first time point of GFP signal appearance could be unambiguously determined were used to generate the scatter plot in Fig 1E.

### Quantitative analysis of TIRF images

For Fig. 1D, image processing and analysis were conducted using ImageJ (NIH). In brief, the following steps were performed: Raw time-lapse images were initially converted to TIFF files. A custom macros program was developed and utilized for the subsequent image analysis (https://github.com/Peii39/Intensity-density-along-the-cilia.git). This program follows a series of systematic steps: 1. The program first reads the input image and allows the user to select a rectangular region as the area of interest. 2. The program will let the user select a segmented line along the cilia, starting from the base to the tip. 3. To enhance the accuracy of the segmented line, the program employs the “Fit Spline” and “Interpolate” functions of ImageJ to achieve a smoother representation. 4. The segmented line is divided into 12 equidistant steps along the cilia length. 5. Each of these 12 segments is enclosed by a polygon with a width parameter (L=250). 6. The program calculates the integrated intensity within each of the 12 polygons. The obtained values are then divided by the individual area size of the respective polygon. The processed data was used to generate an intensity profile with 12 data points along the length of the cilia, illustrating the intensity of GRK2 along the ciliary structure.

For Fig. S2, the image analysis process was conducted using ImageJ (NIH) through a series of distinct steps: 1. Raw time-lapse images were converted into the TIFF file format. 2. The SiR-Tubulin-marked cilia channel movie was imported into Trainable Weka plug-ins to generate a mask based on the SiR-Tubulin signal. In a few of the movies manual delineation was necessary to ensure accuracy in identifying cilia structure. 3. To isolate the region of interest exclusively containing GRK2 within the cilia shaft while excluding the cilia base, a two-step masking process was implemented. First, the “AND” function was applied to the previously generated mask and the rectangular area selected at the cilia base. Subsequently, the “XNOR” operation was employed with the result from the previous step and the original mask to create a new mask that delineated only the cilia shaft area for GRK2 intensity per pixel analysis. 4. Three frames were chosen both before and after the activation of SAG21k, and each set of three frames was subjected to the procedures outlined in steps 1-3 to obtain intensity per unit area under the mask. 5. The background per unit area of these six frames was generated by randomly selecting three circular areas around the cilia mask for background subtraction. 6. The subtraction of the mean of three frames both before and after the activation of SAG21k, following background subtraction, was used to generate a scatter plot. This plot illustrates the mean intensity increase per unit area following SAG21k activation. In a few cases, negative values are observed because of either uneven background signal, or the signal was dim compared to noise or both. In these cases, the precise quantification of signal is not feasible. However, in all 79 cases, signal was clearly observed in the original timelapse images.

### Construction of NIH3T3 *Grk2^-/-^* cell line

CRISPR editing of the endogenous *Grk2* locus was performed via electroporation (Neon, Thermo Fisher) of Alt-R CRISPR reagents (IDT) into NIH3T3 Flp-in cells followed by single-cell FACS sorting of Atto550+ cells, as previously described^38^. The sequence of the sgRNA (Mm.Cas9.GRK2.1.AA) was as follows: /AltR1/rUrC rUrGrG rArArC rArCrG rUrCrC rCrCrU rCrGrG rGrUrU rUrUrA rGrArG rCrUrA rUrGrC rU/AltR2/. Single clones were expanded, lysed in RIPA buffer, and tested for loss of GRK2 via immunoblotting of whole cell lysates. Loss of GLI reporter activation in *Grk2* cells^43,44^ was confirmed using a nano-glo GLI-reporter assay,

### Purification of SMO, GRK2, and PKA-C proteins

FLAG-SMO (SMO residues 64-674) or FLAG-SMOΔCT (SMO residues 64-566, see above) were expressed in HEK293 GnTI^-^ cells, along with GRK2-eGFP, using the BacMam approach in the presence of 10 mM sodium butyrate and 1 µM SAG21k. Cells were collected 48 h after transduction, pelleted, and stored at -80°C for future use. Frozen cell pellets were broken up, and thawed by stirring in a hypotonic lysis buffer (20 mM HEPES pH 7.5, 1 mM EDTA, protease inhibitor tablets, 1 µM SAG21k). The resuspended cells were centrifuged at 20,000 x g at 4°C for 30 minutes and the pellet containing the crude membrane fraction was collected. The crude membrane pellet was broken up by dounce homogenization, then stirred in a solubilization buffer (50 mM HEPES pH7.5, 300 mM NaCl, 5 mM MgCl_2_, 20 mM KCl, 5 mM ATP, 1 mM CaCl_2_, protease inhibitor tablet, 1 µM SAG21k, 1% DDM / 0.1% CHS) at 4°C for 1 h. The solution was then centrifuged at 20,000 x g at 4°C for 30 minutes and the supernatant was subjected to affinity purification using M1 anti-FLAG antibody coupled to Sepharose beads. The column was then washed in 50 mM HEPES pH 7.5, 300 mM NaCl, 1 mM CaCl_2_, 0.025% GDN, and 1 µM SAG21k to exchange 1% DDM, 0.1% CHS with 0.025% GDN and to remove ATP. Protein was eluted with 50 mM HEPES pH 7.5, 150 mM NaCl, 5 mM EDTA, 0.025% GDN, protease inhibitor tablet, 1 µM SAG21k, and 0.2 mg/mL FLAG peptide. For experiments involving non-phosphorylated SMO (Fig. 6C), we omitted the GRK2-eGFP virus during HEK293 expression and utilized 0.25 µM vismodegib in place of SAG21k for all cell growth, purification, and chromatography steps.To limit proteolysis, protease inhibitor tablet was included in the elution buffer and the protein was concentrated and subject to size-exclusion chromatography immediately after affinity purification (within 8h of initially resuspending the frozen cell pellet). For size-exclusion chromatography, the protein sample was concentrated to 500 µL or less and injected onto a Superdex 200 Increase 10/300 GL column equilibrated with 20 mM HEPES, 150 mM NaCl, 0.025% GDN, 1 µM SAG21k. The protein was then injected onto a 500 µL loop on an AktaPure Chromatography workstation. Fractions containing SMO were identified with the FPLC trace from a peak eluting at approximately 11.7 mL, and the fractions were further analyzed via SDS-PAGE analysis. Fractions containing pure SMO were pooled and concentrated. His-tagged GRK2 was expressed in High Five cells via baculovirus and purified via NiNTA affinity chromatography and gel filtration chromatography as previously described^132^. PKA-C was expressed in *E. coli* and purified via IP20 chromatography as previously described^39^.

### Small-scale optimization of GRK2 phosphorylation of SMO *in vitro*

The GRK2 phosphorylation reaction was undertaken using purified agonist- or inverse-agonist-loaded SMO. To obtain inverse-agonist-loaded SMO, SAG21k in a purified SMO-SAG21k complex (see above) was exchanged on resin for the inverse agonist KAADcyc via incubation of SMO-SAG21k with anti-FLAG M2 magnetic beads (Sigma-Aldrich) in a buffer containing 20 mM HEPES pH 7.5, 100 mM NaCl, and 0.05% GDN supplemented with 1 mM KAADcyc at 4°C for two hours. The beads were collected with a magnetic separator and washed five times with 20 mM HEPES pH 7.5, 100 mM NaCl, and 0.05% GDN, along with 100 μM KAADcyc. The protein was eluted with the same wash buffer supplemented with 0.1 mg/ml 3X FLAG peptide.

For *in vitro* kinase assays, 1 μM SMO was first incubated with 2 μM GRK2 and 20 μM c8-PIP_2_ in a buffer containing 20 mM HEPES pH 7.5, 2 mM NaCl, 2 mM MgCl_2_, and 2 mM DTT on ice for 20 minutes, then 50 μM ATP (supplemented with ATPγ^32^P (PerkinElmer) to a final specific activity of 300 Ci/mmol) was added to initiate the reaction. The reaction was incubated at room temperature and quenched by adding SDS sample buffer at various time points. The reaction mixture was separated on SDS-PAGE and the phosphorylated receptor was autoradiographed for image analysis with the Quantity One program (BioRad) on phosphor screens.

For the western blot analysis, the same reaction was set up without radionucleotides. The reaction mixture was separated on SDS-PAGE and then transferred to a nitrocellulose membrane (Bio-Rad) in a transfer buffer (10 mM CAPS pH 11, 10% methanol) using 80 V for 80 minutes. The membrane was blocked with 5% BSA at room temperature for 1 hour and incubated with anti-FLAG M2-peroxidase antibody (Sigma-Aldrich) at room temperature for another hour. After incubation, the blot was washed three times with wash buffer (20 mM Tris pH 7.6, 137 mM NaCl and 0.1% Tween-20), reacted with a chemiluminescent substrate (SeraCare Life Sciences), and imaged using ImageQuant LAS4000.

### In vitro phosphorylation w/GRK2

Purified FLAG-tagged SMO was diluted to 1 µM in a solution containing 20 mM HEPES pH 7.5, 2 mM NaCl, 2 mM MgCl_2_, 100 µM TCEP, 1 µM GRK2, 20 µM PI(4,5)P2 diC8, 0.025% GDN, and 1 µM SAG21k. The solution was left to incubate on ice for 20 minutes before the addition of ATP at a final concentration of 200 µM. The sample was then left to incubate at room temperature for approximately 16h (overnight).

After GRK2 phosphorylation, SMO was separated from GRK2 via FLAG affinity purification using M1 anti-FLAG antibody coupled to Sepharose beads. The column was then washed with 50 mM HEPES pH 7.5, 300 mM NaCl, 1 mM CaCl2, 0.025% GDN, 1 uM SAG21k. The SMO was then eluted with 20 mM HEPES pH 7.5, 150 mM NaCl, 0.2 mg/mL FLAG, 0.025% GDN, 1 uM SAG21k, 5 mM EDTA. Following FLAG affinity purification, SMO was purified further via size exclusion chromatography as described in the SMO purification methods.

### Pulldown assay

SMO and PKA-C were subject to buffer exchange via desalting column into 20 mM HEPES pH 7.5, 150 mM NaCl, 3 mM CaCl_2_, 10 mM MgCl_2_, 1 mM ATP, 0.025% GDN, and 1 µM SAG21k. SMO and PKA-C were combined into a 100 µL solution of the above buffer at concentrations of 18.2 µM and 10 µM respectively. Each binding reaction was applied to 20 µL of sepharose beads coupled to FLAG antibody in a 1.5 mL Eppendorf tube and incubated on a rotator at 4°C for 2h. The flow-through fraction was removed from the beads using a 50 µL Hamilton syringe, then the beads were washed twice with 200 µL of the buffer used above. SMO was then eluted by adding 60 uL of 20 mM HEPES pH 7.5, 150 uL NaCl, 0.2 mg/mL FLAG peptide, 0.025% GDN, 1 µM SAG21k, and 5 mM EDTA, followed by incubation on a rotator at room temperature for 10 minutes. The bead suspension was then centrifuged, the supernatant recovered with a Hamilton syringe, and analyzed on a reducing 4-20% SDS-PAGE gel. Band intensities for PKA-C and SMO on StainFree SDS-PAGE gels (which detect protein via an incorporated UV-activated tryptophan-reactive dye) were quantified via densitometry using ImageLab software, and corrected for the tryptopan content of each protein. The competition binding experiment in Fig. 6D utilized a similar procedure, except that a PKIα(5-24) peptide [TTYADFIASGRTGRRNAIHD, custom-synthesized by PepMic], or a control peptide [EDQVDPRLIDGK], were included at 100 μM in the initial binding mixture, and anti-FLAG M2 magnetic beads were used instead of anti-FLAG M1 sepharose beads.

### Zebrafish forebrain and eye development studies

Embryos were treated with DMSO or SAG21k starting at 3-4 hpf. Small punctures were made in the chorion to ensure accessibility of the small molecule. At 24 hpf, embryos were fixed in 4% PFA in 0.5% PBST at room temperature for 1 hour. Embryos were stained with the following concentrations: anti-pSmo (1:500); anti-acetylated tubulin (1:100); Alexa Fluor 488 Phalloidin (1:250); DAPI (1:1000). Secondary antibodies (Alexa Fluor 568 goat anti-rabbit and Alexa Fluor 647 goat anti-mouse) were used at 1:200. Embryos were mounted in 1.6% low melting point agarose and cleared in glycerol for imaging. Z-stacks were acquired on a Zeiss LSM 980.

### Zebrafish spinal cord studies

Embryos were fixed at 24 hpf in 4% PFA overnight at 4°C. Embryos were stored in 4% PFA for up to 1 week. Immunofluorescence was performed as previously described with several modifications (Tay et al., 2013). Briefly, embryos were blocked in 1X PBS with 0.5% Triton X-100, 5% goat serum, and 2 mg/mL BSA for 4 hrs at room temperature. 1:500 anti-pSmo and 1:400 anti-acetylated tubulin (Sigma T6793) primary antibodies were added and incubated overnight at 4°C. Primary antibodies were removed and samples were washed with two quick washes and then one overnight wash at 4°C in PBS + 0.5% Triton x-100. Embryos were blocked again for 2 hrs at room temperature before addition of donkey anti-rabbit Alexa fluor 488 and donkey anti-mouse Alexa fluor 568 secondary antibodies at 1:200 and topro3 at 1:500 for 2 hrs at room temperature. Samples were again subject to two quick washes and then one overnight wash as described above. Samples were embedded in low melt agarose on a glass bottom dish and imaged using an Olympus Fluoview FV1200 confocal microscope and Olympus FV10-ASW v4.1 software. An Olympus UPlanF1 100X/1.30 objective was used.

### Mouse neural tube studies

All mice were cared for in accordance with NIH guidelines and Emory University’s Institutional Animal Care and Use Committee. Alleles used were: *Smo^cbb^* [MGI: 5911831], *Smo^bnb^* [MGI: 2137553], and *SmoM2* [MGI: 3576373] which was activated through Cre recombination by CMV-cre [MGI: 2176180]. *Ptch1^-/-^* is the *Ptch1^tm1Mps^* lacZ reporter null allele [MGI: 1857447]^88^. Timed matings were performed to generate embryos of the indicated stage. Embryos were dissected in cold PBS and processed for immunofluorescence. After fixing for 1h at 4°C in 4% PFA (prepared in PBS; Thermo Fisher Scientific, catalog 043368.9M), embryos were rinsed with PBS and then cryoprotected in 30% sucrose in 0.1M phosphate buffer overnight at 4°C. Embryos were rinsed and embedded in OCT, before being sectioned at 10microns.

Slides were blocked with antibody wash buffer (0.1% Triton X-100, 1% goat serum in TBS) for 1h at room temperature. Sections were stained with rabbit anti-pSMO (1:1000) and mouse anti-Arl13b (1:1500) for 1h at room temperature. After several washes, slides were incubated with secondary antibodies (1:500) and Hoechst (1:5000) for 1h at room temperature. Following additional washes, slides were coverslipped with Prolong Gold and allowed to cure overnight at room temperature in the dark.

Slides were imaged on a Nikon A1R confocal microscope. Z stacks were acquired using a 60x oil immersion Plan Apochromat TIRF lens through NIS Elements software (Nikon Instruments, version 5.21.03). Post-acquisition max intensity Z projection and image presentation were performed using FIJI/ImageJ (version 1.53t). To assess pSMO and total SMO staining in the mouse neural tube sections, images were coded to remove genotype and antibody information. Six independent researchers marked the length of specific staining in each image. The average length was divided by the total lumen length to determine the percentage of observed staining.

### Analysis of primary CGNPs

Cerebella from postnatal day 7 (P7) C57BL/6J mice were cut into small pieces and incubated at 37°C for 20min in 15U/ml papain solution and DNase I in Hanks’ Buffer with 20 mM Hepes (HHBS). HHBS was used to rinse tissue once and then removed. Tissues were then triturated in Neurobasal medium containing DNase I to obtain single cell suspension. Cells were centrifuged at 1000 rpm for 5 min at 4°C and resuspended in Neurobasal medium containing B-27 Supplement, GlutaMAX Supplement and 1% Pen Strep. Cells were plated on Poly-D-Lysine containing Laminin coverslips at 1.2 x 10^6^ cell/ml. After 36h in culture, cells were pretreated for 4 h with GRK2 inhibitor (Cmpd101 or 14as at 30 μM, or a vehicle control), and then DMSO or 100 nM SAG was added for 8h, and then cells were fixed in 4% PFA for immunostaining. CGNPs were blocked with blocking buffer (0.2% Triton X-100, 2% Donkey serum in PBS) for 1 h at room temperature. After blocking, cells were incubated with rabbit anti-pSMO (1:1000) or rabbit anti-SMO (1:1000)^31^, along rat anti-ARL13B (1:500) at 4°C overnight. Subsequently, cells were incubated with secondary antibodies (ThermoFisher Scientific) for 1 hour and Hoechst 33342 for 10min at room temperature. Cells were mounted in Fluoromount-G. Imaging was done on a Nikon spinning disc confocal system with 100X oil immersion lens.

### Ethics statement

All animal experiments were approved by the respective Institutional Animal Care and Use Committees, as follows: 1) Institutional Animal Care and Use Committee, University of California, Merced, Protocol #: AUP 21-0002. (primary mouse cerebellar granule neuronal precursor studies); 2) Emory University Institutional Animal Care and Use Committee, Protocol # 201700587 (mouse neural tube studies); 3) University of Utah Institutional Animal Care and Use Committee, Protocol # 21-09017 (zebrafish spinal cord studies); 4) University of Utah Institutional Animal Care and Use Committee, Protocol # 21-01007 (zebrafish brain and eye studies).

